# Fibroblast TGF-β3 promotes tissue-residency and survival of CD8 T cells in barrier tissues and tumors

**DOI:** 10.64898/2026.05.20.726599

**Authors:** Sunny Z. Wu, Ryan S. Lane, Alessandra Castiglioni, Endi K. Santosa, Alberto Guarnieri, Apple Cortez Vollmers, Christian Cox, Yagai Yang, Hannah Bender, Tianhe Sun, Justin A. Shyer, Akshay T. Krishnamurty, Sören Müller, Shannon J. Turley

## Abstract

Fibroblasts are key organizers of tissue architecture and immune cell homeostasis, yet how they shape adaptive immune function within non-lymphoid tissues remains incompletely understood. CD8^+^ tissue-resident memory T cells (T_RM_) provide localized protection against pathogens and contribute to tumor control, but the microenvironmental signals that maintain their persistence and survival are poorly defined. Here, we identify fibroblast-derived TGF-β3 as a conserved stromal niche factor that specifically sustains CD8^+^ T_RM_ in both steady-state and disease settings. Across human single-cell cross-tissue atlases, CD8^+^ T_RM_ preferentially correlated with fibroblast abundance in healthy barrier tissues and multiple tumor types, and *TGFB3* emerged as a key fibroblast-enriched candidate mediator. In human and murine co-culture systems, fibroblast-derived TGF-β3 promoted CD8 T_RM_-like differentiation in vitro. Using a novel genetic in vivo model, inducible fibroblast-specific deletion of *Tgfb3* reduced CD8 T_RM_ across barrier tissues at steady state and impaired antigen-specific CD8 T_RM_ formation following viral infection. In tumor models, genetic loss or antibody mediated neutralization of TGF-β3 impaired CD8 T cell residency and cytotoxicity, induced dysfunction via proteotoxic stress and apoptotic programs, and accelerated tumor growth. These findings provide mechanistic insight into the limited efficacy of pan-TGF-β blockade in cancer therapy. Collectively, we describe a novel fibroblast-CD8 T cell axis mediated by TGF-β3 that sustains residency and restrains proteotoxic stress in barrier tissues and tumors.

## Introduction

Fibroblasts are central organizers of tissue architecture and immune homeostasis, shaping the extracellular matrix (ECM) and establishing localized immune niches through dynamic paracrine signaling. In solid tumors, cancer-associated fibroblasts (CAFs) constitute a dominant stromal compartment that governs immune exclusion, therapeutic resistance, and responsiveness to immune checkpoint blockade^1–3^. Fibroblasts regulate CD8 T cells in lymphoid organs^4–6^ and inflamed tissues^1,7,8^, yet whether fibroblast-derived signals actively sustain CD8 T cell survival, function and persistence in non-lymphoid tissues, following infection and in tumors, remain incompletely defined.

Memory T cells are key mediators of adaptive immunity that circulate through blood, secondary lymphoid organs and non-lymphoid tissues, adopting distinct functional states. CD8 tissue-resident memory T cells (T_RM_) represent a specialized differentiation state that persists long term within tissues without recirculating, providing rapid, localized protection against pathogens^9^. In viral infections, CD8 T_RM_ are defined by cell-surface markers including CD103 (*ITGAE*), CD69, CXCR6 and CD49a^10,11^, and transcriptional programs governed by HOBIT, BLIMP1^12^ and RUNX3^13^, among others. In addition to their role in pathogen defense, CD8 T_RM_ have emerged as a prognostically favorable population strongly associated with improved survival and enhanced immunotherapy responses in cancer^14–17^, and experimental models demonstrated a direct contribution of CD8 T_RM_ to tumor control^18^. In tumors, exhausted CD8 T cells (T_EX_) can share transcriptional and phenotypic features with CD8 T_RM_^19,20^, suggesting that tissue-residency and dysfunction may be governed by shared microenvironmental constraints.

A defining feature of CD8 T_RM_ is their dependence on local microenvironmental signals for differentiation and persistence. Genetic ablation of TGF-β-receptor signalling (TGF-βR2) in CD8 T cells impairs T_RM_ establishment in tissues, demonstrating a central requirement for local TGF-β cues^10,21^. Importantly, TGF-β signaling has also been implicated in maintaining stem-like CD8 T cells in tumors^3^ and chronic infection^22,23^, positioning this pathway at the intersection of tissue retention, exhaustion, and therapeutic response. In tumors, TGF-β signaling contributes to immune exclusion^2^ and drives the activation of CAFs^1^. Accordingly, combining TGF-β pathway inhibition with checkpoint blockade enhances antitumor immunity in preclinical models^2,24,25^, partly by expanding stem-like CD8 T cells^3^. However, pan-neutralization of TGF-β has been limited by systemic toxicities and lack of clinical efficacy^26^, underscoring the need to resolve isoform- and cell type-specific functions of TGF-β signaling during homeostasis and in cancer.

TGF-β comprises three isoforms (TGF-β1, TGF-β2 and TGF-β3) with distinct context and cellular expression patterns^27^. TGF-β1 is broadly expressed, whereas TGF-β3 expression is more restricted and enriched within stromal compartments, including fibroblasts. Despite binding the same canonical receptor complex, whether TGF-β isoforms exert divergent, or even opposing, functions within the TME remains unclear.

CD8 T_RM_ exhibit substantial phenotypic and functional diversity across non-lymphoid tissues^28–32^, suggesting that multiple local cues sustain tissue-residency. The requirement for TGF-β signaling is highly tissue-specific^28–31^, with skin and intestinal T_RM_ depending on TGF-β, whereas liver T_RM_ form independently of this pathway^30^. In tumors, this complexity is intensified by stromal remodeling, hypoxia, and chronic antigen exposure, conditions that impose proteotoxic and metabolic stress on infiltrating CD8 T cells^33^. These pressures promote dysfunctional or exhausted states that transcriptionally overlap with T_RM_ populations^19,20^.

Although TGF-β has classically been attributed to epithelial sources for CD8 T_RM_ maintenance, CD8 T_RM_ also provide their own source of TGF-β in an autocrine manner^34^, and naïve CD8 T cells are preconditioned in lymph nodes via dendritic cell-mediated TGF-β activation^35^. Consistent with this context dependency, CD8 T_RM_ precursors transit through deeper layers of tissue, such as the crypt of the intestine^21^ and the dermis of the skin^36^, before establishing epithelial residence, raising the possibility that CD8 T cells receive instructive signals from stromal cells within these regions. Whether stromal cells deliver signals such as TGF-β that shape CD8 T cell persistence in tissues and tumors remains unresolved.

Building on these insights, we hypothesized that fibroblasts provide distinct signals that sustain tissue-resident CD8 T cell survival and function in tissues and tumors. TGF-β3 emerged as a fibroblast-enriched candidate regulator of these states. We combined computational analysis of human single-cell atlases with in vitro co-culture systems, fibroblast-restricted genetic deletion models, and isoform-specific neutralization strategies to define the role of TGF-β3 in regulating CD8 T_RM_ maintenance, survival, and antitumor immunity.

## Results

### CD8+ T_RM_ associate with fibroblast abundance in barrier tissues and tumors

Whilst CD8 T_RM_ heterogeneity has been characterized in mouse tissues^28–31^, the stromal determinants that sustain CD8 T_RM_ maintenance in human tissues and tumors remain poorly defined^37^. To identify microenvironments enriched for CD8 T_RM_, we queried SCimilarity^38^, a cell foundation model trained on >24 million single-cell RNA sequencing (scRNA-seq) profiles, using reference CD8 T_RM_ signatures derived from human lung^39^, breast cancer^14^, and ulcerative colitis^40^ datasets (Fig. 1a). Cells with high similarity to annotated CD8 T_RM_ were further refined using a core CD8 TRM gene signature^10^ relative to circulating T cells (Fig. 1b-c; Extended Data Fig. 1a). This approach identified 643,231 CD8 T_RM_ across 21 tissue types and 13 disease indications (Fig. 1d). CD8 T_RM_ were enriched in multiple barrier tissues, including lung, skin, and gut, and were present across both healthy and disease settings.

**Figure 1.**
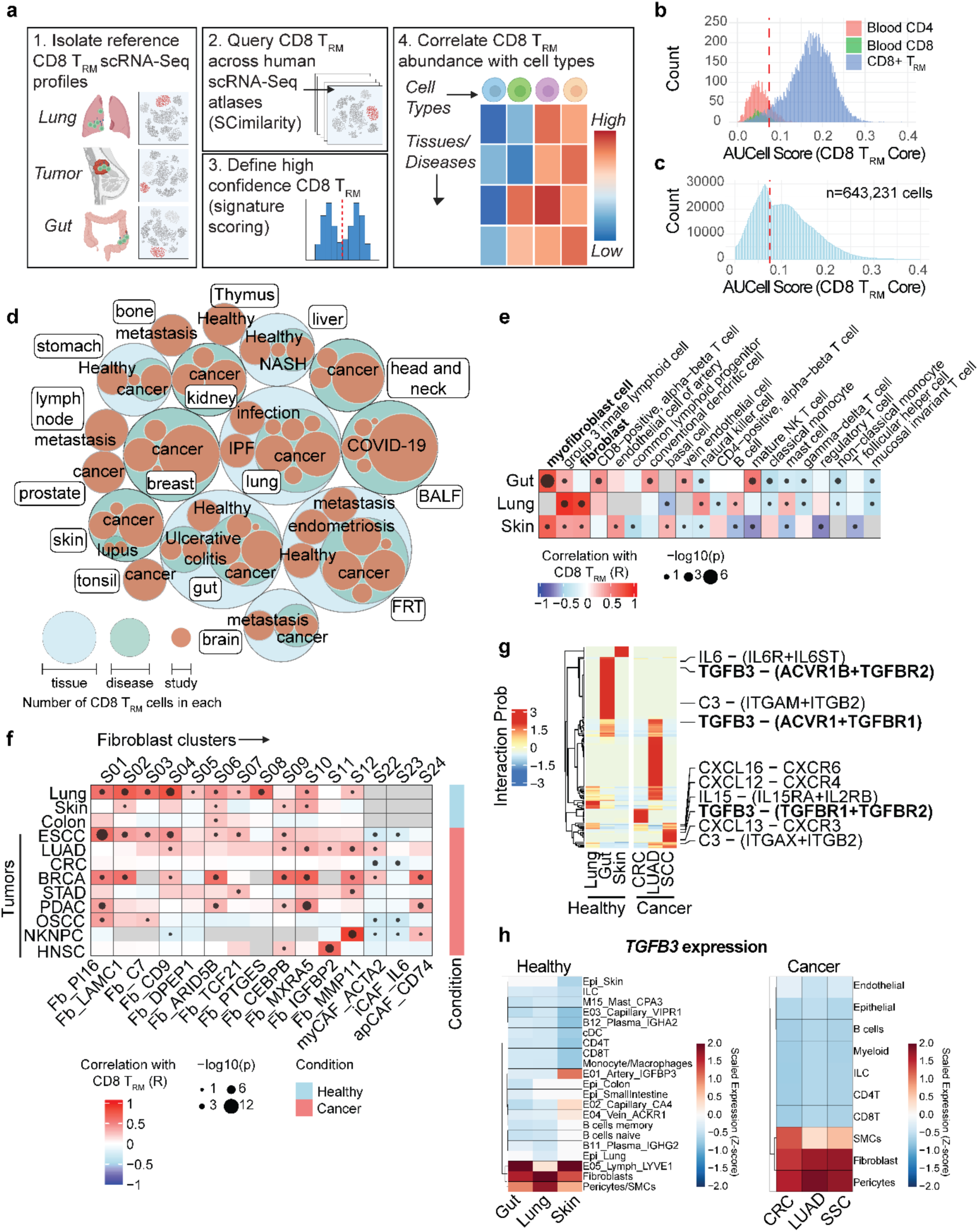
CD8+ T_RM_ correlate with fibroblast abundance in barrier tissues and tumors via *TGFB3* interactions. **a,** Schematic of the computational workflow used to identify CD8⁺ tissue-resident memory T (T_RM_) cells across publicly available scRNA-Seq datasets. The SCimilarity cell foundation model was used for cell query using CD8 T_RM_ from breast, lung and gut tissue studies as reference. **b-c**, Histograms of independent AUCell signature scores using the core CD8 T_RM_ from Mackay et al. 2016^12^. The AUCell threshold (indicated by the red line) for additional filtering was determined using the reference CD8 T_RM_ against circulating T cells (b). Threshold was then applied to all cells identified using SCimilarity (c). **d,** Bubble plot summary of all CD8 T_RM_ cells detected using SCimilarity. Circle size denotes the number of cells detected within each tissue, disease state, and individual study. **e**, Heatmap of pearson correlation analysis between the frequency of CD8⁺ T_RM_ cells and the frequency of major cell populations across gut-, lung- and skin-derived datasets. Cell types were defined using SCimilarity-based annotation. **f**, Heatmap of pearson correlation analysis between independently annotated Itga1^+^ CD8⁺ T_RM_ and fibroblast clusters in the cross-tissue atlas from Shi et al 2025^41^. For e and f, the size of each point represents statistical significance calculated using -log_10_(p-value) and boxes in grey represent insufficient numbers of cells or samples for correlation analysis. **g**, Cellchat cell-cell interaction analysis between fibroblasts and CD8⁺ T_RM_ clusters in barrier tissue-related healthy and tumors datasets. Top interactions ranked by significance in Cellchat are highlighted. **h,** *TGFB3* expression across annotated cell types in barrier tissue-related healthy and tumors datasets. Expression is shown as normalized transcript levels scaled per cell type. For e and f, statistical significance was determined using two-sided Pearson correlation testing (P < 0.05).

To define cellular correlates of CD8 T_RM_ abundance, we performed cell-cell association using Pearson correlation analyses within each tissue context. Across lung, skin, and gut datasets, fibroblasts consistently ranked among the strongest positively correlated cell populations with CD8 T_RM_ (Fig. 1e). Notably, strong associations were observed between CD8 T_RM_ and fibroblasts in ulcerative colitis (R = 0.79), lung infection (R = 0.87), and SLE/DLE skin (R = 0.70), suggesting a conserved stromal-CD8 T_RM_ relationship across barrier tissues.

We next validated these findings in an independent cross-tissue single-cell atlas encompassing healthy and cancer samples^41^. An independently annotated CD8 T_RM_ cluster (‘CD8T03_Trm_ITGA1’) exhibited significant positive correlations with multiple fibroblast states across healthy barrier tissues and tumors, with PI16^+^, ARID5B^+^, CEBPB^+^ and MMP11^+^ clusters among the most frequently associated subsets, including in lung adenocarcinoma (LUAD) and esophageal squamous cell carcinoma (ESCC) (Fig. 1f). In contrast, CAF states (‘S23_myCAF_ACTA2’ and ‘S22_iCAF_IL6’) were negatively correlated with CD8 T_RM_ abundance across several tumor types, consistent with their reported immunosuppressive roles^1^. Notably, these fibroblast associations were not broadly observed with total CD8 T cell infiltration, indicating that the relationship is preferentially enriched for CD8 T_RM_ populations (Extended Data Fig. 1c).

To identify candidate mediators of fibroblast-CD8 T_RM_ crosstalk, we performed ligand-receptor inference using CellChat^42^ across barrier tissues. TGF-β3-associated interactions emerged among the most consistently enriched pathways linking fibroblasts to CD8 T_RM_, alongside chemokine and cytokine signaling (Fig. 1g). Examination of ligand expression revealed that *TGFB3* was predominantly expressed by mesenchymal cells including fibroblasts with minimal expression in immune or epithelial compartments (Fig. 1h). Pericytes and SMCs also expressed *TGFB3*, however, did not exhibit significant correlation with CD8 T_RM_ (Fig. 1e, Extended Data Fig. 1d). Together, these analyses identify fibroblasts as a conserved cellular correlate of CD8 T_RM_ and illuminate fibroblast-derived TGF-β3 as a candidate paracrine regulator of CD8 T_RM_ maintenance in barrier tissues and tumors.

### Fibroblast-derived TGF-β3 promotes a T_RM_-like CD8 T cell state in vitro

Having identified fibroblasts as a conserved correlate of CD8 T_RM_ abundance across barrier tissues and tumors, we next directly tested whether fibroblast-derived signals directly instruct a CD8 T_RM_-like state. Primary human lung fibroblasts were co-cultured with activated human CD8 T cells in a transwell system to isolate contact-independent paracrine effects without confounding allogeneic responses (Fig. 2a). Across independent donor combinations, fibroblasts induced robust numerical expansion of activated CD45RO^+^CD45RA^-^ CD8 T cells that acquired a T_RM_-like phenotype defined by CD103 and CD69 co-expression (Fig. 2b-c).

**Figure 2.**
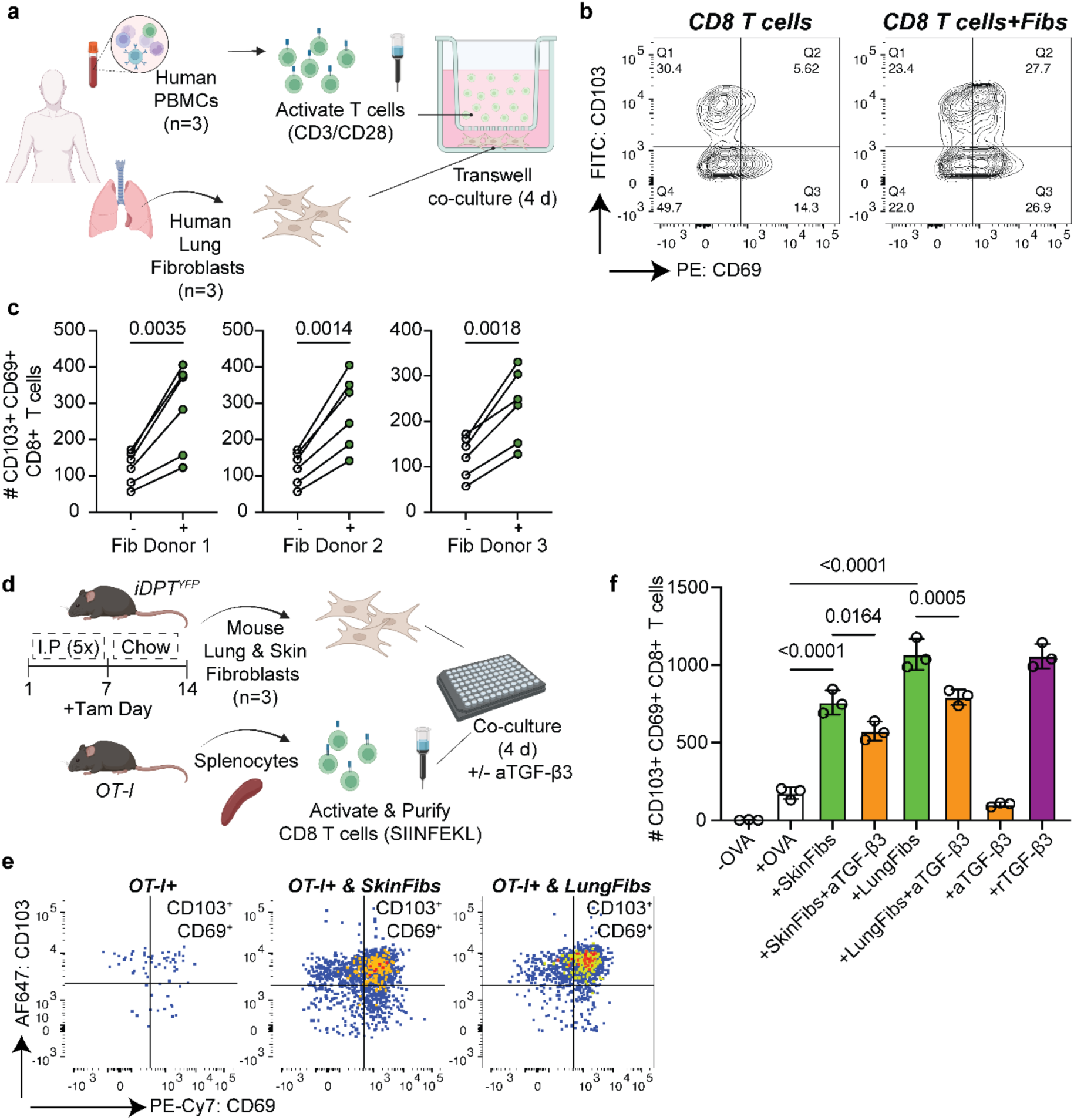
Human and mouse fibroblasts promote a CD8 T_RM_-like state in vitro via TGF-β3. **a**, Schematic of the human peripheral blood mononuclear cell (PBMC)-lung fibroblast co-culture assay performed in a transwell system to assess fibroblast-derived soluble factors. **b**, Representative flow cytometry plots showing induction of a T_RM_-like phenotype in activated CD45RO^+^CD45RA^-^ CD8^+^ T cells, defined by CD103 and CD69 expression. Mono-culture activated CD8 T cells (left) and lung fibroblast co-culture (right) conditions are shown. **c**, Quantification of activated CD103^+^CD69^+^ CD8⁺ T_RM_-like cells in mono-culture versus lung fibroblast co-culture conditions. Each dot represents an individual PBMC donor (n = 3 donors pooled from two independent experiments). Fibroblasts from three independent lung donors were used. Statistical significance was determined using a two-sided paired t-test. **d**, Schematic of the murine OT-I T cell-fibroblast co-culture assay used to test the functional role of TGF-β3 in vitro. **e**, Representative flow cytometry plots showing fibroblast induction of a T_RM_-like phenotype in activated TCRβ⁺CD8⁺CD62L^LOW^CD44^HIGH^ OT-I T cells, defined by CD103 and CD69 expression. OT-I T cells treated with OVA (left), and in co-culture with skin (middle) or lung fibroblasts (right) are shown. **f**, Quantification of CD103⁺CD69⁺ CD8⁺ T_RM_-like OT-I cells following mono-culture, co-culture with lung- or skin-derived fibroblasts, in the presence of an anti-TGF-β3 neutralizing antibody or treatment with recombinant TGF-β3. Each dot represents an independent fibroblast biological replicate. Representative data from four independent experiments are shown. Statistical significance was determined using a two-sided unpaired t-test.

Given the established requirement for TGF-β signaling in T_RM_ differentiation and its predicted interaction with barrier tissue fibroblasts, we focused on TGF-β3 as a candidate mediator. To directly test TGF-β3-specific function, we established a complementary mouse co-culture system using *Dpt*^+^ lung or skin fibroblasts isolated from *iDpt^YFP^* reporter mice^43^ and antigen-specific OT-I T cells (Fig. 2d). Consistent with the human system, mouse fibroblasts promoted a numerical expansion of activated CD62L^LOW^CD44^HIGH^ OT-I T cells that upregulated CD103 and CD69 (Fig. 2e-f). Importantly, neutralization using a TGF-β3-specific neutralizing monoclonal antibody (2A10)^44^ significantly attenuated fibroblast-induced CD103^+^CD69^+^ T_RM_-like differentiation (Fig. 2f), demonstrating a functional requirement for fibroblast-derived TGF-β3. Conversely, recombinant TGF-β3 alone was sufficient to induce a CD8 T_RM_-like phenotype in the absence of fibroblasts (Fig. 2f). Although partial inhibition following TGF-β3 blockade suggests additional fibroblast-derived signals contribute to T_RM_ programming, these data establish TGF-β3 as a paracrine factor driving fibroblast-mediated induction of a CD8 T_RM_-like state.

### Inducible fibroblast-specific deletion of *Tgfb3* enables in vivo dissection of paracrine function

To directly test whether provision of TGF-β3 by pan-tissue *Dpt*+ fibroblasts is required to maintain CD8 T_RM_ in vivo, a new inducible conditional knockout mouse model was generated by crossing *Dpt^IRESCreERT2^;Rosa26^LSLYFP^* mice to *Tgfb3^fl/fl^* mice^45^ containing a *Tgfb3* floxed allele with LoxP sites flanking exon 6, encoding the majority of the mature forms of TGF-β proteins (hereafter referred to as *iDpt^YFP^Tgfb3^fl/fl^;* Fig. 3a). Cre recombination was induced by intraperitoneal tamoxifen administration once daily for five consecutive days, followed by continuous maintenance on tamoxifen-containing chow for the duration of the study (days 14-15) (Fig. 3b). Efficient deletion of *Tgfb3* in *Dpt*⁺ fibroblasts was confirmed by bulk RNA sequencing (Extended Data Fig. 2a-c).

**Figure 3.**
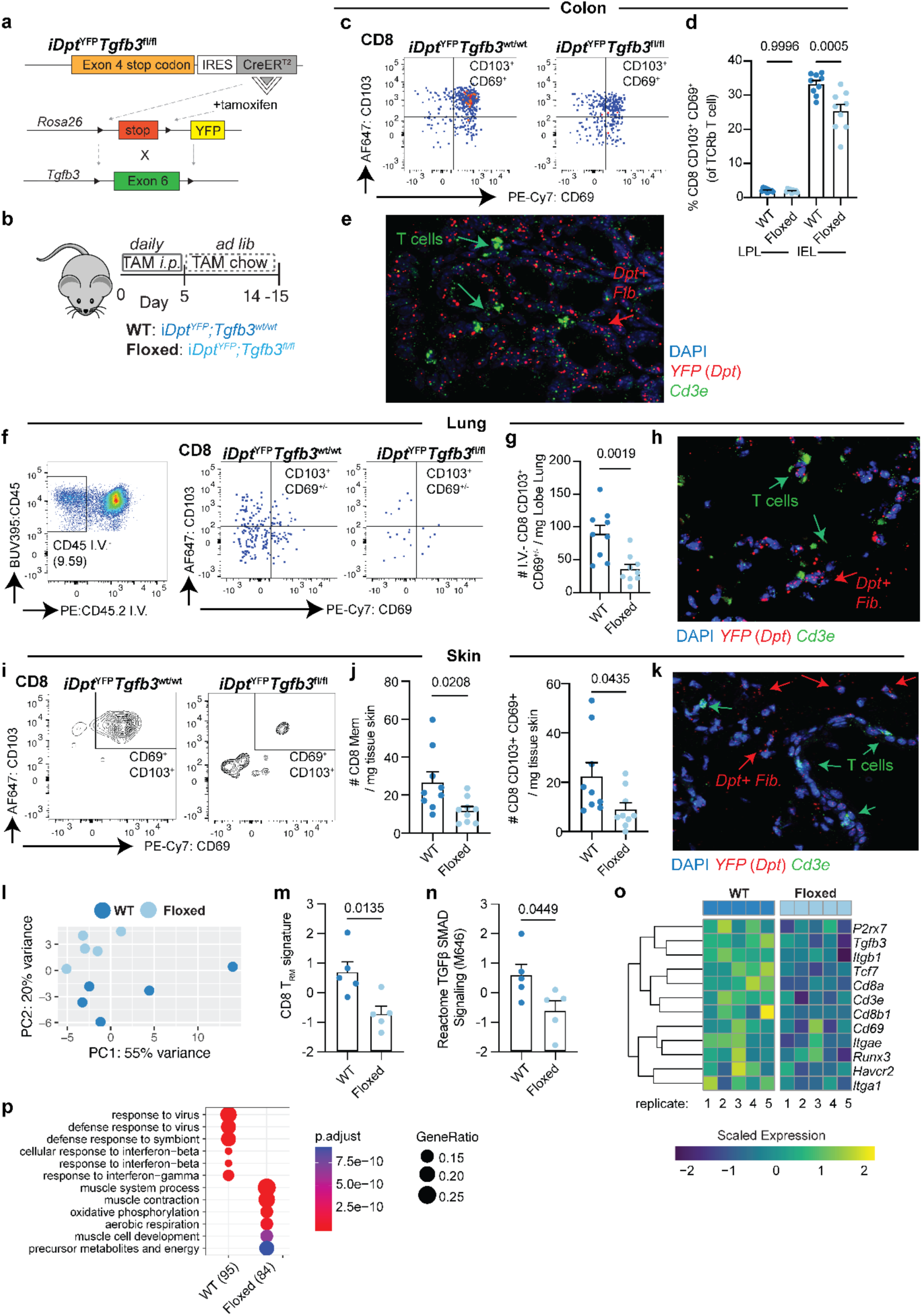
Deletion of *Dpt*+ fibroblast derived *Tgfb3* reduces barrier tissue CD8 T_RM_ in vivo. **a,b**, Schematic of the genetic strategy (a) and experimental workflow (b) used to generate inducible *Dpt^IRESCreERT2^;Rosa26^LSLYFP^* fibroblast-specific *Tgfb3*-deficient mice (*iDpt^YFP^Tgfb3^fl/fl^*). **c**, Representative flow cytometry plots showing colonic CD8^+^ T_RM_, defined as CD45^+^TCRβ^+^ CD44^HIGH^CD62L^LOW^ CD103^+^CD69^+^. Colon compartments were analyzed separately as intraepithelial lymphocytes (IEL) and lamina propria lymphocytes (LPL). **d,** Quantification of the frequency of colonic CD8^+^ T_RM_ in *iDpt^YFP^Tgfb3^fl/fl^*mice compared to controls (of TCRβ^+^ cells). **e**, Representative images from ISH staining of *YFP* (red) and *Cd3e (*green) in the colon of *iDpt^YFP^Tgfb3^wt/wt^* control mice. **f**, Representative flow cytometry plots of lung CD8^+^ T_RM_ defined as intravenously labeled CD45I.V.^-^CD45^+^TCRβ^+^CD44^HIGH^CD62L^LOW^CD103^+^ CD69^+/-^. **g**, Quantification of lung CD8^+^ T_RM_ in *iDpt^YFP^Tgfb3^fl/fl^*mice compared to control animals as total cell numbers per mg of lung lobe. Representative images from ISH staining of *YFP* (red) and *Cd3e* (green) in lungs of *iDpt^YFP^Tgfb3^wt/wt^* control mice. **i**, Representative flow cytometry plots of skin CD8⁺ T_RM_ cells, defined as CD45^+^TCRβ^+^CD44^HIGH^ CD62L^LOW^ CD103^+^CD69^+^. **j**, Quantification of skin CD44^HIGH^CD62L^LOW^ CD8^+^ and CD8⁺ T_RM_ in *iDpt^YFP^Tgfb3^fl/fl^* mice compared to controls. Data are shown as total cell numbers per mg of skin tissue. **k**, Representative images from ISH staining of *YFP* (red) and *Cd3e* (green) in skin of *iDpt^YFP^Tgfb3^wt/wt^*control mice. **l,** PCA of whole-skin bulk RNA-Seq data from *iDpt^YFP^Tgfb3^fl/fl^*and controls. **m-n**, Mean scoring of a core CD8^+^ T_RM_ transcriptional program^12^ (m) and TGF-β / SMAD Reactome (n). **o,** Heatmap of representative DEGs identified by DESeq2. **p**, GO pathway analysis of the top 200 DEGs (top six per genotype are shown). For e, h and k, representative images shown from 6 biological replicates per tissue. Statistical significance was determined using two-way ANOVA in d and two-sided unpaired Student’s t-tests in g, j, m and n.

Enumeration of fibroblast frequency, abundance, global transcriptional programs, and histological analyses across barrier tissues were unchanged (Extended Data Fig. 3a-g), indicating that TGF-β3 is dispensable for fibroblast maintenance or identity. Thus, this model enables selective interrogation of the paracrine role of fibroblast-derived TGF-β3 on immune populations without confounding alterations in the stromal compartment.

### Fibroblast-derived TGF-β3 sustains barrier tissue T_RM_ in vivo

We next assessed immune compartments in *iDpt^YFP^Tgfb3^fl/fl^* mice to determine whether fibroblast-specific *Tgfb3* deletion selectively impacts tissue-resident T cells. Lymph node cellularity and transcriptional profiles were unchanged relative to controls (Extended Data Fig. 4), consistent with the low abundance of *Dpt*^+^ fibroblasts in secondary lymphoid organs^43^.

Similarly, macrophage (F4/80^+^) and dendritic cell (MHC-II^+^CD11c^+^) populations were unaffected across skin, lung, and colon (Extended Data Fig. 5a-e), indicating preserved myeloid homeostasis.

In contrast, fibroblast-specific *Tgfb3* deletion resulted in a selective reduction of tissue-resident T cells across barrier tissues. In the colon, CD8 T_RM_ within the intraepithelial lymphocyte compartment (IEL) were reduced by approximately ∼1.4- and 2-fold in both frequency and absolute number, respectively (Fig. 3c-d; Extended Data Fig. 6a-b). To contextualize these intestinal findings spatially, we next examined *Tgfb3* expression in a published mouse gut spatial transcriptomics dataset^21^. This revealed an enrichment of *Tgfb3* expression at the base of intestinal crypts and within the lamina propria, co-localizing with fibroblast (*Pdgfra*) and CD8 T cell (*Cd8a*) transcripts (Extended Data Fig. 7a). Consistent with this spatial relationship, in situ hybridization (ISH) confirmed localization of *YFP*^+^*/Dpt*^+^ fibroblasts within *Cd3e*^+^ T cell-enriched stromal niches along the crypt-villus axis (Fig. 3e; Extended Data Fig. 7b), supporting a model in which fibroblast-derived TGF-β3 sustains T_RM_ along this colon axis.

A similar phenotype was observed in the lung, where intravascular labeling distinguished resident from circulating T cells (Fig. 3f). Lung CD8 T_RM_ were numerically reduced by ∼2.3-fold in *iDpt^YFP^Tgfb3^fl/fl^*mice (Fig. 3f-g; Extended Data Fig. 6e-f). ISH analysis further revealed that *Cd3e*^+^ T cells localized within *YFP*^+^*/Dpt*^+^ fibroblast-rich adventitial niches in the lung (Fig. 3h; Extended Data Fig. 7c). In steady-state skin, CD44^HIGH^ CD62L^LOW^ CD8 T cells and CD103^+^CD69^+^ T_RM_ were reduced by ∼1.8- and ∼2-fold despite their lower baseline abundance, respectively (Fig. 3i-j; Extended Data Fig. 6j-k). *YFP*^+^*/Dpt*^+^ fibroblasts were broadly distributed throughout the dermal layer of the skin, including stromal niches in proximity to *Cd3e*^+^ T cells (Fig. 3k; Extended Data Fig. 7d). CD4 T cells expressing CD69 were also diminished in colon (Extended Data Fig. 6c-d) and lung (Extended Data Fig. 6h-i), though not skin (Extended Data Fig. 6l), indicating tissue-specific sensitivity of CD4 residency programs to fibroblast-derived TGF-β3. Together, these findings demonstrated that fibroblast-derived TGF-β3 is required for the maintenance of CD8 T_RM_, and in select tissues CD69^+^ CD4 T cells, across multiple barrier organs, while leaving lymphoid tissues and myeloid compartments intact.

### Loss of fibroblast-derived *TGF-β3* reshapes skin immune transcriptional programs

To determine whether the reduction in skin CD8 T_RM_ was reflected at the tissue level, we performed bulk RNA sequencing of steady-state skin following fibroblast-specific *Tgfb3* deletion. PCA revealed modest but consistent transcriptional separation between *iDpt^YFP^Tgfb3^fl/fl^*and control samples (Fig. 3l). Consistent with flow cytometry, *Tgfb3*-deficient skin exhibited reduced enrichment of a CD8 T_RM_ gene signature (Fig. 3m), accompanied by decreased expression of broad T cell markers (*Cd3e, Cd8a, Cd8b1, Tcf7*), T_RM_-associated integrins (*Itgae, Itga1, Itgb1*), and residency-linked genes such as *P2rx7* (Fig. 3o). TGF-β/SMAD pathway activity was also significantly reduced (Fig. 3n), confirming impaired local TGF-β signaling within the skin microenvironment.

Gene ontology and Hallmark pathway analyses further revealed depletion of interferon-α and interferon-γ response programs in *Tgfb3*-deficient skin (Fig. 3p; Extended Data Fig. 6m). Given that skin-resident CD8 T_RM_ constitutively express and respond to interferon-related pathways^32^, these transcriptional changes are consistent with loss of resident CD8 T cell-driven immune gene expression. Overall, these data demonstrated that fibroblast-derived TGF-β3 maintains tissue-resident CD8 T cells and associated interferon programs within barrier tissues, reinforcing its role as a stromal regulator of local immune function.

### Fibroblast-derived *TGF-β3* is required for virus-specific CD8 T_RM_ establishment

To determine whether fibroblast-derived TGF-β3 is required for the formation of antigen-specific CD8 T_RM_ following acute infection, we employed a localized Modified Vaccinia virus Ankara (MVA) ear infection model^46^ (Fig. 4a). Tamoxifen-treated *iDpt^YFP^Tgfb3^fl/fl^* and control mice were infected via ear scarification, and virus-specific CD8 T cells were tracked at day 7 and day 30 post infection using H-2K(b) tetramers loaded with the immunodominant B8R peptide (TSYKFESV) (Fig. 4b).

**Figure 4.**
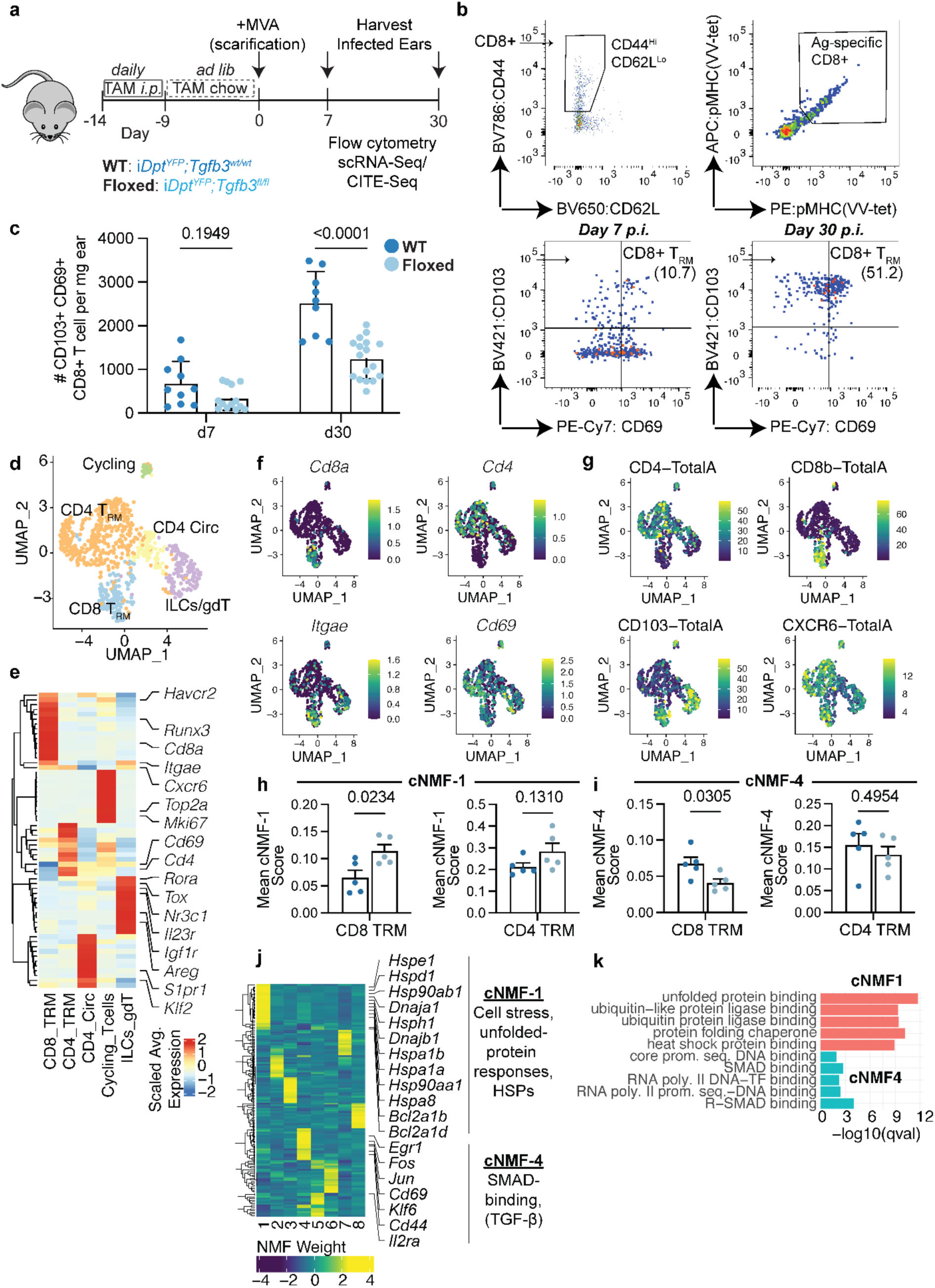
Fibroblast-specific *Tgfb3* deficiency impairs virus-specific CD8 T_RM_ formation following localized skin infection. **a**, Experimental schematic of the localized ear infection model using MVA virus in *iDpt^YFP^Tgfb3^fl/fl^* mice. Infected ear tissue was harvested at day 7 and day 30 post infection. **b**, Representative flow cytometry plots of virus-specific CD8^+^ T_RM_ cells identified as CD45^+^ TCRβ^+^ CD44^HIGH^CD62L^LOW^ CD103^+^ CD69^+^ cells binding H-2K(b) tetramers loaded with the vaccinia virus B8 peptide (TSYKFESV). Tetramers conjugated to two independent fluorophores (APC and PE) were used to ensure specificity. Representative plots from day 7 and day 30 post infection are shown. **c**, Quantification of total virus-specific CD8^+^ T_RM_ cells in infected ears of *iDpt^YFP^Tgfb3^fl/fl^*and control mice at day 7 and day 30 post infection. **d**, UMAP visualization of single-cell transcriptomic profiling of T cells isolated from infected ears at day 30 post infection. **e**, Cluster-averaged heatmap of significantly differentially expressed genes across T cell clusters identified in d. **f,g**, Feature plots of canonical T cell and T_RM_ marker genes measured at the RNA level (f) and corresponding protein expression detected by antibody-derived tags (ADT) (g). **h,i,** Quantification of mean biological program scores for consensus non-negative matrix factorization (cNMF) program 1 (h) and program 4 (i) in CD8^+^ and CD4^+^ T_RM_ cells from *iDpt^YFP^Tgfb3^fl/fl^*and control mice. **j**, Heatmap showing gene loadings for eight identified T cell cNMF programs. Highly weighted genes contributing to cNMF-1 and cNMF-4 are highlighted. **k,** Gene ontologies enriched in the top 100 gene loadings weighted for the cNMF-1 and cNMF-4 programs. Statistical significance was determined using two-way ANOVA for c and two-sided unpaired Student’s t-tests for h,i.

At day 30 post infection, when effector CD8 T cells have transitioned into long-lived CD8 T_RM_, *iDpt^YFP^Tgfb3^fl/fl^*mice exhibited a significant reduction in B8R-specific CD44^HIGH^ CD62L^LOW^ CD8 T cells co-expressing CD103 and CD69 within infected skin (Fig. 4c). This defect mirrored the steady-state CD8 T_RM_ loss observed in uninfected skin, indicating that fibroblast-derived TGF-β3 is required not only for T_RM_ maintenance but also for their optimal establishment following infection.

In contrast, virus-specific effector memory, central memory, and naïve CD8 T cell populations in draining cervical lymph nodes at day 30 post infection were unchanged (Extended Data Fig. 5f-g), demonstrating intact priming and differentiation. These data establish a tissue-local requirement for fibroblast-derived TGF-β3 in the formation and persistence of antigen-specific CD8 T_RM_ following acute viral challenge.

### CD8 T_RM_ up-regulate a proteotoxic unfolded protein stress response in the absence of fibroblast *Tgfb3*

To define the transcriptional mechanisms by which loss of fibroblast-derived TGF-β3 impairs CD8 T_RM_ persistence, we performed scRNA-Seq profiling of T cells isolated from infected skin at day 30 post-infection. A total of five infected mice per *iDpt^YFP^Tgfb3^fl/fl^* and *iDpt^YFP^Tgfb3^wt/wt^* group (total n=10) were analyzed. Within the T cell compartment, a total of five clusters were identified. These included CD4 and CD8 T_RM_ that expressed combinations of canonical tissue-resident marker genes *Itgae* (CD103)*, Cd69, Cxcr6, Havcr2* and *Runx3,* and cell surface CD4 or CD8b, CD103 and CD69; Circulating CD4 T cells (CD4_CIRC_) expressing *S1pr1* and *Klf2;* proliferating T cells expressing *Mki67* and cell cycle related genes including *Top2a;* and a cluster of innate lymphoid cells (ILCs) and gamma-delta T cells (ILCs_gdT) defined by markers *Rora, Nr3c1, Il23r* and *Igf1r* (Fig. 4d-g).

To identify biological gene programs in CD8 T_RM_ that depend on fibroblast *Tgfb3*, consensus non-negative matrix factorization (cNMF) was performed across all T cell clusters. A total of eight distinct cNMF programs were resolved, of which two (cNMF-1 and cNMF-4) were significantly altered in CD8 T_RM_ from *iDpt^YFP^Tgfb3^fl/fl^* mice compared to controls (Fig. 4h-j). cNMF-4 was enriched for tissue-residency-associated genes, including *Cd69*, as well as components of TGF-β signaling and downstream SMAD-responsive transcriptional targets, including *Egr1*, *Fos*, *Jun*, and *Cd44*. This program was significantly reduced in CD8 T_RM_ isolated from *iDpt^YFP^Tgfb3^fl/fl^* mice relative to *iDpt^YFP^Tgfb3^wt/wt^* controls (Fig. 4i-k). Diminished TGF-β-associated transcriptional activity is consistent with impaired activation of residency machinery in the absence of fibroblast-derived TGF-β3, paralleling prior studies demonstrating defective T_RM_ formation following TGF-β receptor deficiency^10^. Consistent with steady-state observations in skin, cNMF-4 was not significantly altered in CD4 T_RM_ (Fig. 4i).

In contrast, cNMF-1 was significantly elevated in CD8 T_RM_ from *iDpt^YFP^Tgfb3^fl/fl^* mice (Fig. 4h). Genes with high loading in this program included numerous molecular chaperones and heat-shock proteins, such as *Hspe1, Hspd1, Hsp90ab1, Hsph1, Hspa1b, Hspa1a, Hspa8, Hsp90aa1, Dnaja1, and Dnajb2*, as well as BCL2-family members associated with apoptotic regulation (*Bcl2a1b* and *Bcl2a1d*) (Fig. 4j). Gene ontology analysis of cNMF-1 revealed enrichment for ubiquitin-dependent protein degradation, unfolded protein response (UPR), and chaperone-mediated protein folding pathways, consistent with activation of proteostasis and cellular stress programs (Fig. 4k). Such transcriptional programs associated with dysregulated proteostasis and proteotoxic stress responses (PSR) have recently been linked to T cell dysfunction in chronic infection and cancer^33,47,48^. While not statistically significant, a similar upward trend in the proteostasis-associated cNMF-1 program was observed in CD4 T_RM_ from *iDpt^YFP^Tgfb3^fl/fl^* (Fig. 4h).

Overall, these findings indicate that the reduction in viral-specific CD8 T_RM_ following fibroblast-specific *Tgfb3* deletion is not solely attributable to diminished TGF-β-dependent residency signaling. Rather, loss of fibroblast-derived TGF-β3 induces a proteotoxic stress program in CD8 T_RM_, implicating impaired cellular fitness and survival as a complementary mechanism driving T_RM_ attrition within the tissue microenvironment.

### Isoform-specific TGF-β3 neutralization impedes T cell cytotoxicity and promotes proteotoxic stress response

Given that loss of fibroblast-derived TGF-β3 induced a proteotoxic stress program in viral-specific CD8 T_RM_, we next asked whether this axis similarly regulates CD8 T cells within tumors, where dysregulated proteostasis has been linked to tissue-residency and chronic T cell dysfunction in cancer^33,47,48^. Prior studies using pan-TGF-β-neutralizing antibodies (targeting all three isoforms) demonstrated synergy with anti-PD-L1 (aPD-L1) in preclinical models^3^, suggesting that TGF-β signaling broadly restrains antitumor immunity. However, whether individual TGF-β isoforms exert distinct or opposing functions in the TME remains unclear.

To dissect isoform-specific contributions, we re-examined the immune-excluded EMT6 tumor model^3^ using antibodies selectively neutralizing TGF-β1 or TGF-β3 (2A10)^44^, alongside pan-TGF-β blockade (1D11). Blockade of TGF-β1 in combination with aPD-L1 induced significant tumor regression, outperforming control treatment, single-agent aPD-L1, and pan-TGF-β neutralization (Fig. 5a). Consistent with transcriptional changes following pan-TGF-β neutralization^3^, TGF-β1 blockade combined with aPD-L1 significantly expanded transcriptional signatures associated with CD8 stem-like memory T cells (T_SCM_) and interferon-γ responses (Fig. 5b-c). This was accompanied by induction of genes related to CD8 T cell activation and cytotoxicity, including *Ifng, Ifngr1, Tnf, Il15ra, Gzmk, Gzmb, Gzma, Ccl3, Ccl4, Icos*, and *Ly6c1*, as well as memory-associated genes such as *Ccr2, Cxcr3, Zfp36l2,* and *Gpr183* (Fig. 5d).

**Figure 5.**
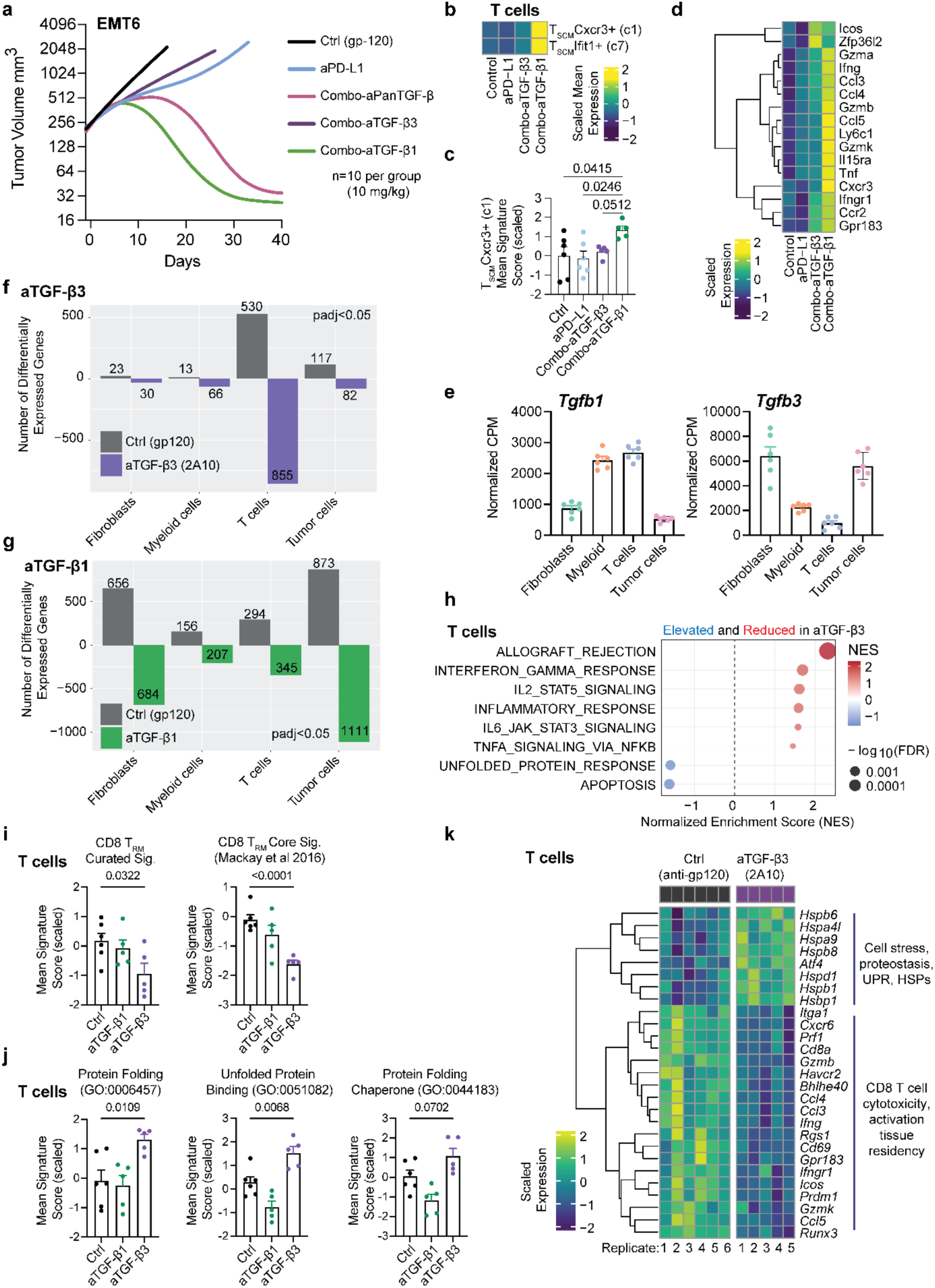
Isoform-specific TGF-β3 neutralization impedes T cell cytotoxicity and promotes proteotoxic stress responses. **a**, Tumor growth kinetics of the EMT6 syngeneic breast tumor model treated with isotype control (gp120), anti-PD-L1 (aPD-L1), aPD-L1 combined with pan-TGF-β neutralization, aPD-L1 combined with anti-TGF-β1, or aPD-L1 combined with anti-TGF-β3 isoform-specific neutralization. **b,c,** Bulk RNA-sequencing analysis of treated EMT6 tumors. Heatmap (b) and quantification (c) of a CD8⁺ memory stem-like gene signature derived from Castiglioni et al., 2023^3^. **d**, Heatmap of selected CD8⁺ T cell cytotoxicity-associated genes, demonstrating preferential induction in tumors treated with the aPD-L1 + anti-TGF-β1 combination. **e**, Expression of *Tgfb1* and *Tgfb3* across cell compartments in control (gp-120) treated EMT6 tumors. Normalized counts per million is shown. **f-g,** Summary of the total number of significant differentially expressed genes (adjusted P < 0.05, DESeq2) across fibroblast, myeloid, T cell, and tumor cell compartments following treatment with anti-TGF-β3 monotherapy (f) or anti-TGF-β1 monotherapy (g) compared with control (gp120) without aPD-L1. **h**, Gene set enrichment analysis (GSEA) of Hallmark pathways differentially regulated in EMT6 tumor-infiltrating T cells following control (gp120) or anti-TGF-β3 treatment. Pathways reduced following anti-TGF-β3 are shown in red; pathways elevated are shown in blue. **i**, Quantification of curated and core CD8^+^ T_RM_ gene signatures derived from Mackay et al., 2016^12^. **j**, Quantification of three proteostasis-related gene ontology (GO) signatures in tumor-infiltrating T cells. **k,** Heatmap of selected differentially expressed genes in T cells from control (gp120) versus aTGF-β3-treated tumors. Statistical significance for c, i, and j was determined using two-sided unpaired Student’s t-tests.

In contrast, blockade of TGF-β3 in combination with aPD-L1 failed to improve tumor control (Fig. 5a) and did not enhance T_SCM_ or interferon-γ transcriptional programs beyond single-agent aPD-L1 (Fig. 5b-d). These findings indicate that TGF-β1 is the dominant immunosuppressive isoform in this tumor model, whereas TGF-β3 does not mediate the same suppressive function and may instead support CD8 T cell function within the TME.

To isolate the specific role of TGF-β3 independent of checkpoint blockade, tumors were treated with anti-TGF-β1 or anti-TGF-β3 monotherapy and compared to isotype control. Fibroblast, myeloid, tumor cell, and T cell compartments were sorted and subjected to bulk RNA sequencing. Analysis of ligand expression revealed that *Tgfb3* was enriched in fibroblasts and tumor cells, whereas *Tgfb1* was predominantly expressed by myeloid and T cell populations (Fig. 5e). Among all cellular compartments analyzed, T cells exhibited the greatest number of differentially expressed genes following TGF-β3 neutralization (n = 1,385; padj < 0.05), indicating that T cells represent the primary population impacted by TGF-β3 blockade (Fig. 5f). In contrast, neutralization of TGF-β1 resulted in broader transcriptional changes across multiple cellular compartments, consistent with its broader expression pattern (Fig. 5g).

Gene set enrichment analysis (GSEA) of T cells from anti-TGF-β3-treated tumors revealed significant depletion of immune activation pathways, including allograft rejection, IFN-γ response, IL2/STAT5 signaling, IL6/JAK/STAT3 signaling, and TNF signaling via NF-κB, consistent with impaired CD8 T cell effector function (Fig. 5h). Notably, these changes were accompanied by enrichment of pathways associated with apoptosis and unfolded protein response (UPR)/proteostasis dysregulation, paralleling the proteotoxic stress signature observed in viral-specific CD8 T_RM_ following conditional *Tgfb3* deletion (Fig. 4k).

Importantly, TGF-β3 neutralization, but not TGF-β1 blockade, resulted in significant depletion of two independent CD8 T_RM_ gene signatures, including a manually curated signature and a core CD8 T_RM_ signature derived from Mackay et al. 2016^12^. This included reduced expression of tissue-residency-associated genes *Cxcr6, Cd69, Runx3, Rsg1, Bhlhe40, Itga1* and *Prdm1*, as well as activation and cytotoxic mediators *Ifng*, *Ifngr1, Gzmb*, *Gzmk, Tnf, Ccl3, Ccl4, Ccl5* (Fig. 5i,k). These findings mirror the phenotype observed in the fibroblast-specific *Tgfb3* conditional knockout model, further supporting a conserved role for TGF-β3 in sustaining CD8 T_RM_ programs in tissues and tumors.

Consistent with this loss of residency-associated transcription, TGF-β3 blockade induced enrichment of multiple proteotoxic stress response signatures, including protein folding (GO:0006457), unfolded protein binding (GO:0051082), and protein folding chaperones (GO:0044183). These programs included induction of *Atf4* and numerous molecular chaperones (*Hspa4l, Hspa9, Hspb8, Hspb1, Hsbp1, Hspd1, Hspb6*) (Fig. 5j-k), further implicating disrupted proteostasis and cellular stress as a consequence of TGF-β3 neutralization.

Collectively, these findings reveal isoform-specific functions of TGF-β signaling in tumors, in which TGF-β1 acts as a dominant immunosuppressive mediator restraining checkpoint responsiveness, whereas TGF-β3 supports CD8 T cell tissue-residency, cytotoxic fitness, and proteostatic stability. Thus, TGF-β3 exerts a protective role in maintaining CD8 T cell function in tumors, paralleling its requirement for steady-state and infection-induced CD8 T_RM_ maintenance in barrier tissues.

### Accelerated subcutaneous tumor growth in the absence of fibroblast-derived *TGF-β3*

Given that isoform-specific antibody neutralization of TGF-β3 impaired CD8 T cell survival, cytotoxicity and induced proteotoxic stress programs in tumors, we next asked whether selective genetic deletion of fibroblast-derived *Tgfb3* would similarly compromise antitumor immunity in vivo. Whereas systemic antibody blockade targets all cellular sources of TGF-β3, our conditional model enables fibroblast-restricted deletion, thereby allowing resolution of cell type-specific contributions within the TME. To test this, we utilized a subcutaneous tumor model on a C57BL/6 background using our conditional *iDpt^YFP^Tgfb3^fl/fl^*mice.

The KPR pancreatic ductal adenocarcinoma (PDAC) model was selected due to its high stromal content and prominent infiltration of cancer-associated fibroblasts (CAFs), the majority of which arise from *Dpt*⁺ fibroblasts (KPR; Kras^LSL.G12D/wt^; p16/p19^fl/wt^; p53^LSL.R270H/wt^; Pdx1.Cre)^1^. To enable tracking of endogenous tumor-specific CD8 T cell responses, KPR cells were engineered using a lentiviral piggyBac transposon system to express full-length ovalbumin (KPR^OVA^), allowing detection of OVA-specific CD8 T cells using H-2K(b) tetramers loaded with SIINFEKL peptide.

*iDpt^YFP^Tgfb3^fl/fl^* and *iDpt^YFP^Tgfb3^wt/wt^* control mice were treated with the same tamoxifen regimen described above prior to subcutaneous implantation of KPR^OVA^ tumor cells (Fig. 6a). Tumor growth was monitored for 35-36 days post-implantation. Fibroblast-specific deletion of *Tgfb3* resulted in significantly accelerated tumor progression, reflected by increased tumor volume and tumor weight at day 35-36 in *iDpt^YFP^Tgfb3^fl/fl^* mice compared to controls (Fig. 6b-e). These findings further support a tumor-protective role for fibroblast-derived TGF-β3.

**Figure 6.**
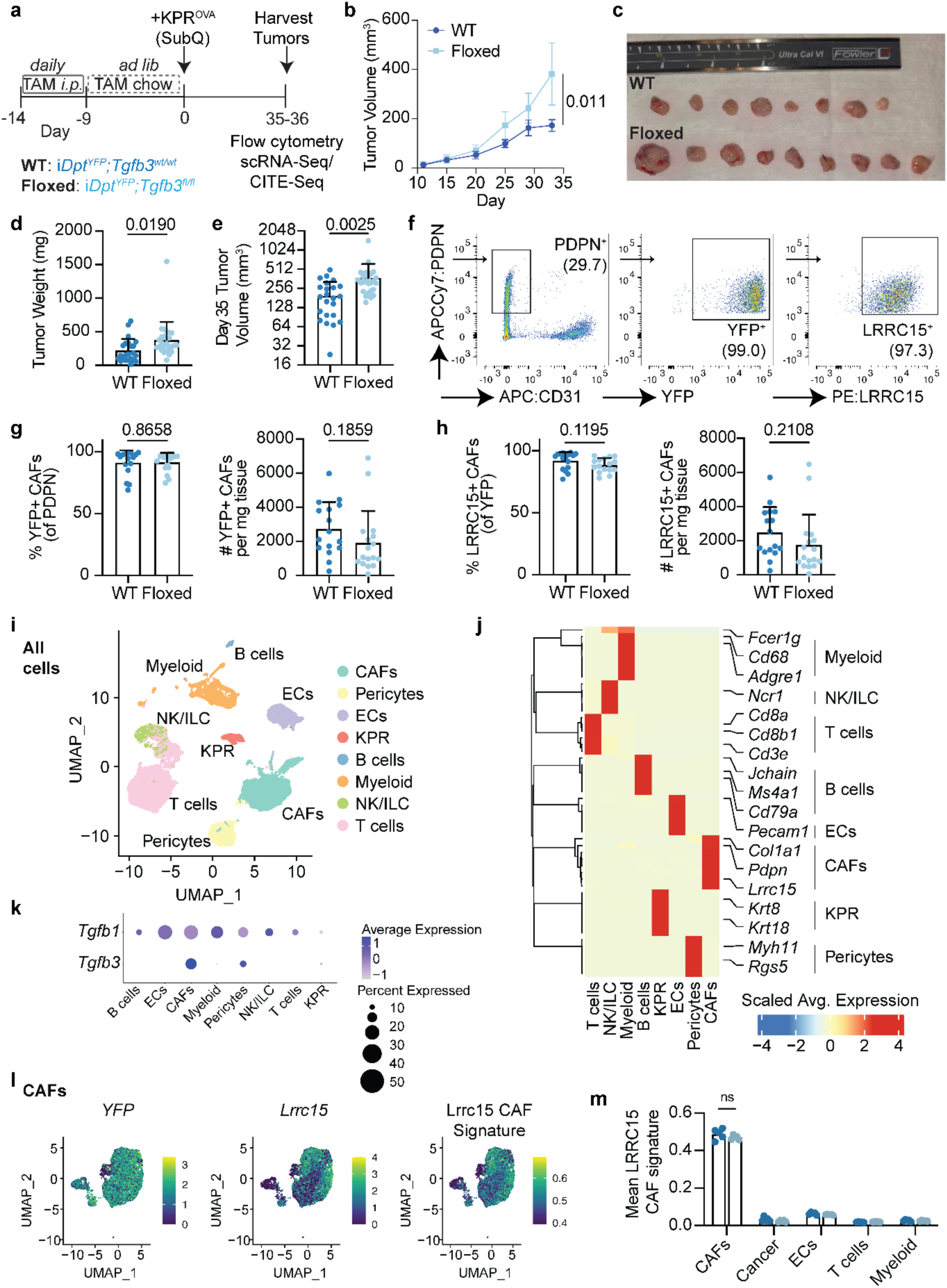
Tumor growth is accelerated in the absence of fibroblast-derived *TGF-β3* in a CAF-independent manner. **a**, Experimental schematic of the KPR^OVA^ tumor model implanted into *iDpt^YFP^Tgfb3^fl/fl^* mice. Mice received tamoxifen intraperitoneally (i.p.) five times over one week, followed by maintenance on tamoxifen-containing chow prior to tumor implantation. Tumors and tumor-draining lymph nodes (tdLNs) were harvested at day 35-36 post implantation. **b,c**, Tumor growth kinetics (b) and representative images (c) of KPR^OVA^ tumors grown in *iDpt^YFP^Tgfb3^fl/fl^* or control mice. Representative data from three independent experiments are shown. **d,e**, Quantification of tumor weights (d) and volumes (e) at endpoint. Data are pooled from three independent experiments. **f**, Representative flow cytometry plots identifying CAFs defined as Live CD24^-^ EPCAM^-^ CD45^-^ CD31^-^ PDPN⁺ cells, with assessment of YFP expression (*Dpt*-lineage tracing) and LRRC15 expression. **g,h**, Quantification of the frequency (left) and total number (right) of YFP⁺ and LRRC15⁺ CAFs in tumors from *iDpt^YFP^Tgfb3^fl/fl^* and control mice. Data are pooled from two independent experiments. **i**, UMAP visualization of single-cell transcriptomic profiling of all cells isolated from KPR^OVA^ tumors at day 36 post implantation. **j**, Cluster-averaged heatmap of significantly differentially expressed genes across major lineage clusters identified in **i**. Selected lineage-defining marker genes are highlighted. **k**, Bubble plot showing *Tgfb1* and *Tgfb3* expression across major tumor lineage clusters. **l**, Feature plot visualization of *YFP* expression, *Lrrc15* expression, and a LRRC15⁺ CAF gene signature score following reclustering of CAF populations. **m**, Quantification of the LRRC15⁺ CAF gene signature in CAFs and other cell types isolated from KPR^OVA^ tumors grown in *iDpt^YFP^Tgfb3^fl/fl^* or control mice. Statistical significance was determined using two-sided unpaired Student’s t-tests for b, d-e, g-h, and two-way ANOVA for m.

### Deficiency in fibroblast-derived *TGF-β3* does not impair CAF or myeloid activation

To determine whether accelerated tumor growth was driven by alterations in stromal or myeloid compartments, tumors were harvested for flow cytometric and single-cell analyses. Given prior reports that TGF-β receptor signaling is required for activation of LRRC15⁺ CAFs, we first assessed whether deletion of *Tgfb3* altered CAF abundance or differentiation through autocrine mechanisms. At 35-36 days post-implantation, approximately 90% of PDPN^+^ fibroblasts in tumors were YFP^+^ and LRRC15^+^, consistent with prior studies demonstrating that the majority of LRRC15^+^ CAFs derive from *Dpt*^+^ fibroblasts^1^. No differences in the frequency or absolute number of total YFP^+^ fibroblasts or LRRC15^+^ CAFs were observed between KPR^OVA^ tumors grown in *iDpt^YFP^Tgfb3^fl/fl^*and *iDpt^YFP^Tgfb3^wt/wt^* controls (Fig. 6f-h).

To further evaluate whether *Tgfb3* deletion altered CAF transcriptional identity, scRNA-seq was performed on tumors from an independent cohort (n = 5 per genotype). Unsupervised clustering identified major cell populations including myeloid cells (*Fcer1g, Cd68, Adgre1*), NK and innate lymphoid cells (*Ncr1*), T cells (*Cd3e, Cd8a, Cd8b1*), B cells (*Jchain, Ms4a1, Cd79a*), endothelial cells (*Pecam1*), CAFs (*Col1a1, Pdpn, Lrrc15*), KPR^OVA^ tumor cells (*Krt8, Krt18*), and pericytes (*Myh11, Rgs5*) (Fig. 6i-j). Across all compartments, fibroblasts and to a lesser extent a small pericyte subset, represented the predominant source of *Tgfb3* expression, whereas *Tgfb1* was broadly expressed across stromal and immune populations (Fig. 6k).

Reclustering of CAFs resolved eight transcriptionally distinct subsets that were largely unchanged in cell neighborhood abundance between *iDpt^YFP^Tgfb3^fl/fl^*and control tumors, as determined by Milo^49^(Extended Data Fig. 8a-c). Quantification of LRRC15^+^ CAF signature scores using AUCell similarly revealed no significant differences between genotypes (Fig. 6l-m), confirming preserved CAF activation states.

Consistent with steady-state tissues, no significant differences were detected in total CD11b^+^ myeloid cells, CD11b^+^F4/80^+^ macrophages, or CD11b^+^F4/80^-^CD11c^+^MHC-II^+^ dendritic cells in tumors or tumor-draining lymph nodes (tdLNs) from *iDpt^YFP^Tgfb3^fl/fl^*mice compared to controls (Extended Data Fig. 9a-f). At single-cell resolution, no coherent genotype-dependent shifts in cell neighborhood abundance were observed across eleven myeloid clusters that mostly included tumor-associated macrophage, monocyte, or DC states (Extended Data Fig. 9g-h).

These data demonstrated that accelerated tumor growth following fibroblast-specific *Tgfb3* deletion occurs independently of changes in CAF abundance or activation state, or myeloid composition. Together with observations in steady-state tissues and antibody neutralization models, these findings support a predominantly paracrine role for fibroblast-derived TGF-β3 in regulating T cell function rather than stromal or myeloid activation.

### Fibroblast-derived *TGF-β3* sustains tumor-specific CD8 T cell cytotoxicity and prevents proteotoxic stress

Having excluded major alterations in CAF and myeloid compartments, we next examined whether impaired CD8 T cell states are altered in tumors of*Dpt^YFP^Tgfb3^fl/fl^* mice compared with controls. Flow cytometric analysis revealed a significant ∼1.7-fold and ∼1.9-fold decrease in the frequency and total number, respectively, of tumor antigen-specific CD44^HIGH^CD62L^LOW^ effector CD8 T cells in *iDpt^YFP^Tgfb3^fl/fl^*tumors compared to littermate controls, tracked using dual-fluorophore H-2K(b) SIINFEKL tetramers (Fig. 7a-b; Extended Data Fig. 10a-c). This reduction was accompanied by decreased CD103^+^ and CD69^+^ tumor-specific CD8 T cells (∼1.3-1.7-fold; Fig. 7c-d), indicating loss of T_RM_-like tumor-infiltrating CD8 T cells within the TME. In contrast, OVA-specific effector memory, central memory, and naïve CD8 T cells in tdLNs were unchanged (Extended Data Fig. 10f-g). Thus, fibroblast-derived TGF-β3 is required locally within the TME to sustain tumor-specific CD8 T cells.

**Figure 7.**
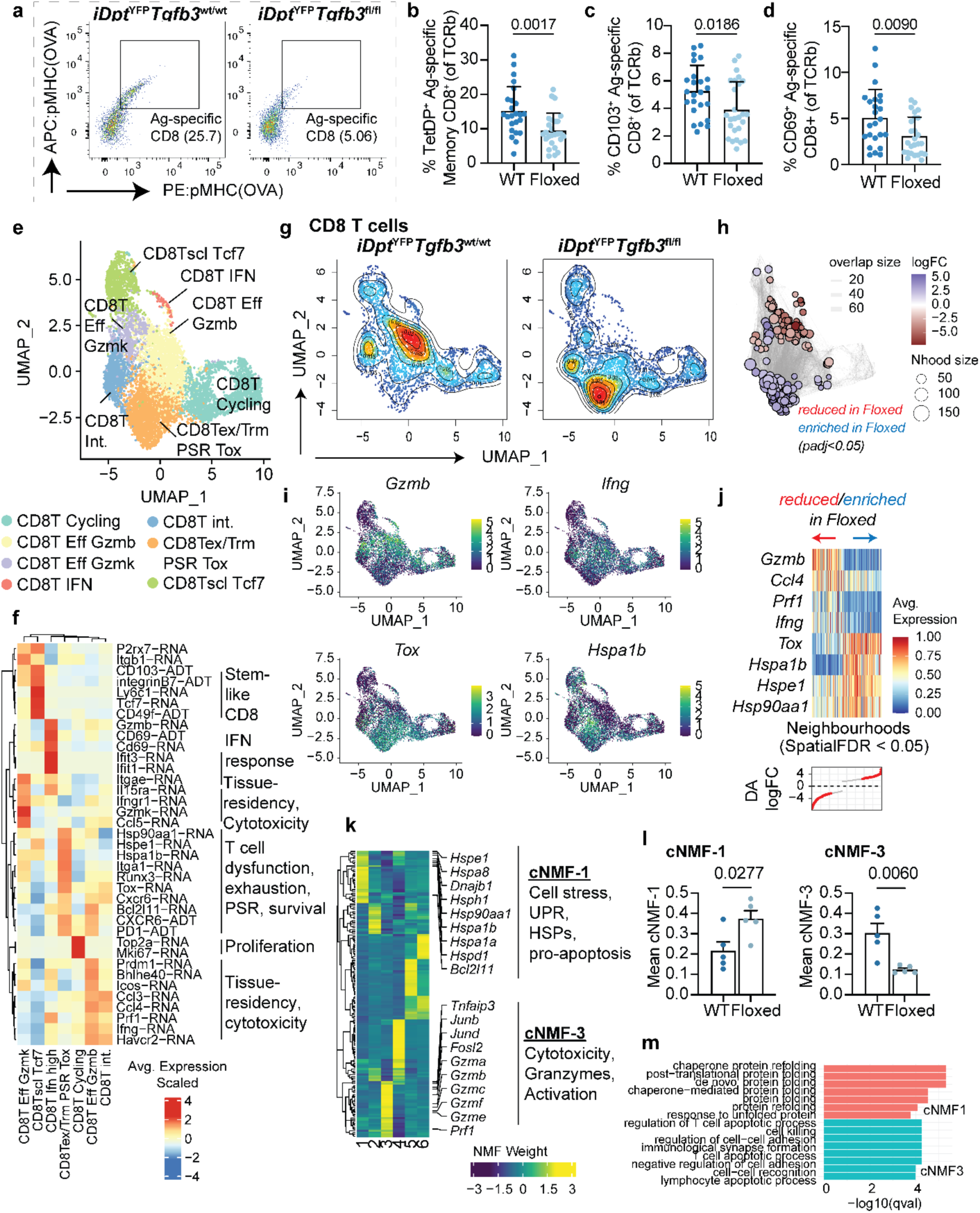
Tumor-specific CD8 T cells are reduced in the absence of fibroblast-derived *TGF-β3* and exhibit dysfunction and proteotoxic stress programs. **a**, Representative flow cytometry plots of OVA-specific CD8⁺ T cells identified as CD45^+^ TCRβ^+^ CD44^HIGH^CD62L^LOW^ cells binding H-2K(b) tetramers loaded with the OVA peptide SIINFEKL. Tetramers conjugated to two independent fluorophores (APC and PE) were used to ensure specificity. Representative plots from KPROVA tumors grown in *iDpt^YFP^Tgfb3^fl/fl^*or control mice are shown. **b-d**, Quantification of total OVA-specific CD8^+^ T cells (b) and subsets expressing CD103 and CD69 (c,d) in subcutaneous KPR^OVA^ tumors harvested at day 35-36 post implantation from *iDpt^YFP^Tgfb3^fl/fl^* and control mice. **e**, UMAP visualization of single-cell transcriptomic profiling of tumor-infiltrating CD8⁺ T cells isolated from KPR^OVA^ tumors at day 36 post implantation. **f**, Cluster-averaged heatmap of significantly differentially expressed genes and surface proteins (antibody-derived tags; indicated as “-ADT”) across major CD8⁺ T cell states identified in e. Selected state-defining markers are highlighted. **g,** UMAP density plots showing the distribution of CD8^+^ T cells derived from control mice (left) or *iDpt^YFP^Tgfb3^fl/fl^* mice (right). **h,** Differential abundance analysis of CD8⁺ T cell neighborhoods using Milo. Neighborhoods significantly reduced in *iDpt^YFP^Tgfb3^fl/fl^* tumors are shown in red, and enriched neighborhoods in blue. **i**, Feature plots showing expression of *Gzmb, Ifng, Tox* and *Hspa1b* across CD8⁺ T cell states. **j,** Neighborhood-averaged expression of selected marker genes demonstrating depletion of *Gzmb*^+^ cytotoxic effector CD8⁺ T cells (*Ccl4, Prf1, Ifng*) and enrichment of *Tox*⁺PD-1⁺ dysfunctional CD8⁺ T cells expressing proteotoxic stress response genes (*Hspa1b, Hspe1, Hsp90aa1*) in *iDpt^YFP^Tgfb3^fl/fl^*tumors. **k**, Heatmap displaying gene loadings for six identified CD8⁺ T cell cNMF programs. Highly weighted genes contributing to cNMF-1 and cNMF-3 are highlighted. **l**, Quantification of mean biological program scores for cNMF-1 (left) and cNMF-3 (right) in CD8⁺ T cells from *iDpt^YFP^Tgfb3^fl/fl^* and control mice. **m**, Gene ontology (GO) enrichment analysis of the top 100 weighted genes from cNMF-1 and cNMF-3 programs. Statistical significance for b-d and l was determined using two-sided unpaired Student’s t-tests.

To define transcriptional changes accompanying this defect, tumor-infiltrating CD8 T cells were profiled by scRNA-Seq. Unsupervised clustering identified seven CD8 T cell states, including stem-like (*Tcf7, Ly6c1*), IFN-responsive, two cytotoxic effector states (*Gzmb*^+^ and *Gzmk*^+^), a transitional population, cycling cells, and a *Tox*^+^PD-1^+^ dysfunctional population enriched for proteotoxic stress response genes (*Hspe1, Hspa1b, Hsp90aa1, Bcl2l11*) (Fig. 7e). Residency-associated markers were broadly expressed across multiple clusters at both transcript and protein levels (Fig. 7f).

Unlike CAF and myeloid compartments, CD8 T cells exhibited pronounced genotype-dependent shifts in state distribution (Fig. 7g). Differential abundance analysis identified significant remodeling of CD8 T cell neighborhoods in *iDpt^YFP^Tgfb3^fl/fl^* tumors (∼21% altered; Fig. 7h). Cytotoxic *Gzmb*^+^ effector neighborhoods expressing *Ccl4, Prf1*, and *Ifng* were selectively depleted, whereas *Tox*^+^PD-1^+^ dysfunctional neighborhoods enriched for proteotoxic stress genes were expanded (Fig. 7h-j).

To resolve underlying gene programs, cNMF was performed. Two programs were significantly altered. A cytotoxic/TGF-β-associated program (cNMF-3), enriched for *Gzma, Gzmb, Gzmk, Prf1* and SMAD-responsive genes (*Junb, Jund, Fosl2, Tnfaip3*), was reduced ∼3-fold in *Tgfb3*-deficient tumors (Fig. 7k-l). In contrast, a proteostasis/UPR-associated program (cNMF-1), enriched for heat-shock proteins and chaperones (*Hspe1, Hspa8, Hsp90aa1, Hspd1*) and the pro-apoptotic gene *Bcl2l11* (Bim), was elevated ∼2-fold (Fig. 7k-m).

Collectively, these data demonstrated that fibroblast-derived TGF-β3 maintains tumor-specific CD8 T cell cytotoxic and residency-associated programs while restraining proteotoxic stress and apoptotic signaling. In its absence, CD8 T cells shift from functional effector states toward stress-associated dysfunctional phenotypes, consistent with the accelerated tumor growth in *iDpt^YFP^Tgfb3^fl/fl^* mice.

## Discussion

Our study highlights the importance of elucidating homeostatic stromal mechanisms in steady-state tissues, as these networks establish immune setpoints that are later co-opted or disrupted in disease. These findings build on mounting evidence that the generation of durable CD8 T_RM_ pools requires sequential, spatially compartmentalized boosts of TGF-β along their activation and migratory trajectories^21,34,35^. This trajectory initiates as early as the naïve T cell stage in the lymph node, where migratory DCs transactivate TGF-β to precondition resting naïve CD8 T cells for epithelial resident fate^35^. Keratinocytes in skin activate sources of latent TGF-β that can be produced by CD8 T_RM_ themselves^34^. We extend this developmental continuum into the barrier tissue stroma, demonstrating that fibroblast-derived TGF-β3 provides a critical survival niche such as in the colon lamina propria and skin dermis. This aligns with recent findings that stromal chemokines^21^ promote CD8 T cell migration into tissues. Together, these data suggest that fibroblast-derived TGF-β3 contributes to this stromal transit and survival phase, reinforcing emerging evidence that T_RM_ interact with neighboring stromal cells to establish local tissue immunity^50^.

Although CAFs are generally characterized as tumor-promoting and immune-suppressive in solid cancers, our findings reveal a previously underappreciated immune-supportive function of the tumor stroma. Prior studies demonstrated that fibroblast TGF-β response signatures correlate with CD8 T cell exclusion and resistance to PD-L1 blockade^2,3^. Specifically, TGF-β receptor signaling in fibroblasts drives the emergence of highly suppressive LRRC15^+^ CAFs that physically and functionally impede CD8+ T cell anti-tumor immunity^1^. In contrast, our genetic and functional analyses indicate that stromal TGF-β signaling is not uniformly immunosuppressive but context-dependent, with fibroblast-derived TGF-β3 rather supporting CD8 T cell survival and tissue-residency and having no impact on fibroblasts themselves. These distinctions may also help explain the clinical limitations of pan-TGF-β blockade. Broad neutralization strategies have shown limited efficacy and dose-limiting toxicities^26^, including cardiac valvulopathies linked to disruption of homeostatic TGF-β2/β3 signaling. Our findings provide an immunological framework consistent with these observations, suggesting that selective targeting of TGF-β1 while preserving TGF-β3 dependent stromal niches may enhance antitumor immunity without compromising T cell survival cues or tissue homeostasis.

The divergent effects of TGF-β ligands remain puzzling, particularly given that TGF-β1 and TGF-β3 isoforms signal through the same type II receptor and canonical SMAD2/3 pathways. While our data indicate that fibroblast-derived TGF-β3 sustains T_RM_ survival, it remains unclear whether this reflects quantitative differences in SMAD signaling, engagement of non-canonical pathways^51–53^, or modulation of TCR signal propagation^54^. An additional layer of regulation likely resides in extracellular control of ligand activation and presentation. All TGF-β isoforms are secreted as latent complexes bound to latency-associated peptides (LAP), and isoform-specific effects may be shaped by differences in matrix association, dependence on integrin-mediated activation, or spatial confinement within stromal niches. Future work defining how TGF-β3 bioavailability is regulated within tissue microenvironments will be essential to understanding how T_RM_ survival niches are maintained.

Our findings also place fibroblast-derived TGF-β3 at the intersection of stromal niche factors and CD8 T cell proteostasis. The TME imposes metabolic stress, nutrient limitation, and chronic antigen exposure, conditions that activate the unfolded protein response (UPR) and contribute to T cell dysfunction^33,47,48^. In contrast, T_RM_ maintain a poised state, utilizing integrated stress response pathways to preserve effector potential while avoiding terminal exhaustion^55^. We propose that fibroblast-derived TGF-β3 functions as an extrinsic regulator of this stress equilibrium, restraining proteotoxic stress responses while preserving cytotoxicity. In its absence, CD8 T cells exhibit increased proteostasis dysregulation and apoptotic signaling, linking a disruption in stromal niche factors to diminished T cell function.

Collectively, our study reframes fibroblasts not solely as architects of immune exclusion in tumors, but as context-dependent regulators of T cell survival and function. By identifying fibroblast-derived TGF-β3 as a stromal factor that sustains CD8 T_RM_ persistence in barrier tissues, these findings highlight the importance of preserving homeostatic stromal cues when modulating TGF-β signaling in cancer. A deeper understanding of how stromal niches calibrate T cell proteostasis and residency may inform strategies to enhance antitumor immunity without broadly disrupting tissue immune homeostasis.

## Methods

### Mice

All animal experimental protocols in this study were approved by the Institutional Animal Care and Use Committee at Genentech. *Dpt^IresCreERT2ki/ki^Rosa26^LSLYFP^* mice^43^ and *Tgfb3^fl/fl^*mice^45^ were designed, generated and bred at Genentech. *Rosa26^LSLYFP^* mice were licensed from Columbia University Irving Medical Center^56^. OTI.Tcr mice were bred and housed at Genentech. Female BALB/c mice were obtained from Charles River Laboratories (Hollister, CA). Female mice aged 6-12 weeks old were used for all experimental studies. Mice were maintained under specific pathogen-free conditions using the guidelines of the US National Institutes of Health.

Sample sizes were based on the number of mice routinely needed to establish statistical significance. Mice were housed in specific pathogen-free, individually ventilated cages following the guidelines of the US National Institutes of Health. Animal rooms were maintained on a 14:10 hour light-dark cycle and were temperature- and humidity-controlled at 68-79°F and 30-70% humidity, respectively, with 10 to 15 room air exchanges per hour. No statistical methods were used to predetermine sample size.

### Generation of iDpt^YFP^Tgfb3^fl/fl^ mice

The *Tgfb3* conditional knockout *iDpt^YFP^Tgfb3^fl/fl^* mice were generated at Genentech using C57BL/6N ES (C2) cells and established methods. *Tgfb3^fl/fl^* mice^45^ contain two loxP sites flanking exons 6 in the *Tgfb3* allele encoding the majority of sequence in the mature forms of TGFb proteins previously targeted in TGFb3 germline knockout mice^57,58^. *Tgfb3^fl/fl^* mice^45^ were crossed with *Dpt^IresCreERT2ki/ki^Rosa26^LSLYFP^* mice^43^ and maintained on a mixed C57BL/6N and C57BL/6J genetic background. Control *iDpt^YFP^Tgfb3^wt/wt^* mice were littermate controls.

### Tamoxifen administration

For induction of Cre expression using tamoxifen, mice were injected with 2 mg tamoxifen (Sigma, cat# T5648) diluted in sunflower seed oil (Sigma, cat# 88921). Injections were performed 5 times daily over 7 days intraperitoneally before switching to chow containing Tamoxifen for a 7 day maintenance (Envigo, cat# TD130.859).

### Skin, lung and lymph node digestion and preparation

For FACS analysis, fibroblasts and immune cell isolation for scRNA-seq or bulk sequencing, indicated tissues were collected into RPMI media containing 10% FCS. Whole tissues were then transferred into 3 ml of digestion buffer (RPMI containing 2% FBS, 0.1mg/ml DNAse I (Roche, #10104159001), 0.2 m/ml Collagenase P (Roche, #11249002001), and 0.8 mg/ml Dispase (Life Technologies, #1710504). Flank skin tissue was processed by first shaving hair and removing adipose tissue. Incubation of the tissues was performed for 15 min at 37°C then triterated by repeated pipetting with a wide-O pipet tip. The supernatant was removed and passed through a 70 µm filter before another 2-3 ml of digestion buffer was added for another 15 min incubation. All tissues were digested for a total of 3 rounds with the exception of lymph nodes and lung that were digested for a total of 1 and 2 rounds, respectively. Lung samples were lysed with RBC ACK lysis buffer for 3 min at room temp. Cells were centrifuged (300xg, 5 min) and blocked with Fc block (BD Pharmingen; #553142, 1:100).

### Colon lamina propria and intraepithelial digestion and preparation

Colon IEL and LPL tissue compartments were isolated as previously described^59^. Briefly, Colon tissue was isolated and removed of adherent adipose tissue and Peyer’s patches. Intestines were opened longitudinally and shaken vigorously in PBS. Tissues were incubated in 25 ml intestinal intraepithelial lymphocyte (IEL) solution made up of 1× PBS, 2% FBS (ThermoFisher), 10 mM HEPES buffer (ThermoFisher), 1% penicillin/streptomycin (ThermoFisher), 1% l-glutamine (ThermoFisher), 1 mM EDTA (Sigma), and 1 mM dithiothreitol (DTT; Sigma). Intestines were vigorously shaken at 250 r.p.m. for 15 min at 37 °C and then removed from IEL suspension, rinsed in PBS and transferred into 50-ml tubes containing ceramic beads (3× one-quarter-inch, MP Biomedicals) in 25 ml collagenase solution made of 1× RPMI 1640 with 2% FBS (ThermoFisher), 10 mM HEPES buffer (ThermoFisher), 1% penicillin/streptomycin (ThermoFisher), 1% l-glutamine (ThermoFisher), 1 mg ml−1 collagenase A (Sigma) and 1 U ml−1 DNase I (Sigma). Intestines were incubated for 30 min at 37 °C with vigorous shaking (250 r.p.m.). Digested LPL samples were passed through a 100-μm strainer, centrifuged and washed by centrifugation (500xg, 5 min).

### Flow cytometry

Single cell suspensions were washed with FACS buffer and stained sequentially with primary antibodies for 20 min in the dark on ice. For MHC tetramer staining experiments, cells were stained with peptide tetramer reagents at room temperature for 20 min immediately prior to primary antibody staining. Fixed peptide monomers were prepared according to manufacturer’s Flex-T protocol recommendations. Flex-T™ Biotin H-2 K(b) VV Monomer (TSYKFESV) (Biolegend, #280053) and Flex-T™ Biotin H-2 K(b) OVA Monomer (SIINFEKL) (Biolegend, #280051) were used for MVA and KPR^OVA^ related studies, respectively. Just prior to analysis, cells were resuspended in MACS buffer containing live dead dye 7-AAD at 1:50 (Biolegend, #420403). Calcein violet staining (Thermo Fisher, #C34858) was additionally used for all fibroblast panels at 1:1000. Cells were analyzed using the BD Symphony (HTS; BD Biosciences) and analyzed via FlowJo (v10.8.1 or v10.9.0).

### Human fibroblast PBMC co-cultures

Human lung fibroblasts were commercially sourced from Lonza from three individual donors. Lung fibroblasts were grown on tissue culture plates coated with rat-tail collagen type I (Sigma, #C3867-1VL) in complete DMEM media under hypoxic conditions (37 C, 1% O2). Human PBMCs were sourced from StemCellTech. Co-culture assays were performed using 6.5 mm 24-well Transwell plates with 0.4 μm pore polycarbonate membrane insert (Corning, #CLS3413), allowing the investigation of secreted factors implicated in fibroblast crosstalk in an allogeneic independent manner. A total of 10,000 human lung fibroblasts were seeded in each well of a collagen-coated 24-well plate a day prior to co-culture. PBMCs were thawed, counted and physiologically activated using Dynabeads Human T-Activator CD3/CD28 (Thermo Fisher, #111.32D) according to the manufacturer’s instructions. A total of 1 M PBMCs were activated in RPMI in a 24-well plate that was treated with 25 ul of prepared dynabead solution. PBMCs were mixed by pipetting and incubated at 37 C for 24 hrs. The following day, PBMCs were collected, washed, centrifuged (500xg, 5 min) and counted. For each co-culture interaction, a total 250k PBMCs (1:25 cell ratio) were placed in the insert at a volume of 100 ul in the 24-well Transwell plate containing fibroblasts. Co-cultures were performed in DMEM media and incubated for 4 days under hypoxic conditions (37 C, 1% O2) prior to flow cytometry analysis.

### Mouse fibroblast OTI co-cultures

*Dpt*+ fibroblasts were isolated from skin and lungs of *Dpt^IresCreERT2ki/ki^Rosa26^LSLYFP^* mice^43^ treated with tamoxifen and prepared as single-cell suspensions as described above. Cells were labeled with Fc block and antibodies, and sorted for CD45^-^EpCAM^-^CD31^-^ PDPN^+^YFP^+^ cells by FACS. Fibroblasts were expanded on tissue culture plates coated with rat-tail collagen type I (Sigma, #C3867-1VL) and grown in complete DMEM in hypoxic conditions (37 C, 1% O2). Fibroblasts were cryopreserved (CryoStor CS10, catalog number 07930) to maintain low passage number. For assays, YFP^+^ fibroblasts were thawed from frozen stock and allowed to recover for 1 week, and passaged once before co-culture. In 96-well tissue culture plates coated with collagen, 5,000 YFP^+^ fibroblasts were plated 1 day prior to co-culture.

Splenocytes were isolated from OTI.Tcr mice into RPMI media, crushed and passed through a 70 µm filter. Cells were treated with RBC ACK lysis buffer for 5 min at room temperature and centrifuged (500xg, 5 min). SIINFEKL peptide (OVA 257-264, #AS-60193-1) was used to activate a total of 20 M OTI cells plated into a 6 well tissue culture plate at a concentration of 0.01 ug / ml in complete RPMI media. After overnight incubation at 37 C for 20-24 hrs, splenocytes were collected, washed and centrifuged (500xg, 5 min). T cells were isolated using Pan T Cell Isolation Kit II, mouse (Miltenyi, #130-095-130) according to manufacturer instructions. A total of 125,000 T cells were co-cultured with fibroblasts (1:25 cell ratio) in DMEM media and incubated for 4 days under hypoxic conditions (37 C, 1% O2) prior to flow cytometry analysis. TGF-β3 antibody neutralization was performed using the previously published 2A10 aTGF-β3 antibody^44^. For controls, OTI T cells were supplemented with 10 ng/ml recombinant human/mouse TGF-β3 (StemCellTech, #50-197-6538).

### RNA Extraction

Mouse lymph nodes, skin, lung, colon IEL and LPL tissues were isolated and dissociated as described above. Total RNA extraction was performed on isolated cell suspensions using the RNeasy Micro kit (Qiagen, #74004). Total RNA was quantified with Qubit RNA HS Assay Kit (Thermo Fisher Scientific) and quality was assessed using RNA ScreenTape on TapeStation 4200 (Agilent Technologies). cDNA library was generated from approximately 2 nanograms of total RNA using Smart-Seq V4 Ultra Low Input RNA Kit (Takara). 150 picograms of cDNA were used to make sequencing libraries by Nextera XT DNA Sample Preparation Kit (Illumina). Libraries were quantified with Qubit dsDNA HS Assay Kit (Thermo Fisher Scientific) and the average library size was determined using D1000 ScreenTape on TapeStation 4200 (Agilent Technologies). Libraries were pooled and sequenced on NovaSeq 6000 (Illumina) to generate 30 millions single-end 50-base pair reads for each sample.

### Bulk RNA sequencing data processing

RNA-sequencing data were analyzed using HTSeqGenie in BioConductor^60^ as follows: first, reads with low nucleotide qualities (70% of bases with quality <23) or matches to rRNA and adapter sequences were removed. The remaining reads were aligned to the human reference genome (human: GRCh38.p10, mouse: GRCm38.p5) using GSNAP^61,62^ 70,71 version ‘2013-10-10-v2’, allowing maximum of two mismatches per 75 base sequence (parameters: ‘-M 2 -n 10 -B 2 -i 1 -N 1 -w 200000 -E 1 --pairmax-rna=200000 --clip-overlap’). Transcript annotation was based on the Gencode genes database (human: GENCODE 27, mouse: GENCODE M15). To quantify gene expression levels, the number of reads mapping unambiguously to the exons of each gene was calculated. 4.0.x: RNA-sequencing data were analyzed using HTSeqGenie in BioConductor as follows: first, reads with low nucleotide qualities (70% of bases with quality <23) or matches to ribosomal RNA and adapter sequences were removed. The remaining reads were aligned to the human reference genome (NCBI Build 38) using GSNAP70,71 version ‘2013-10-10’, allowing maximum of two mismatches per 75 base sequence (parameters: ‘-M 2 -n 10 -B 2 -i 1 -N 1 -w 200000 -E 1 --pairmax-rna=200000 --clip-overlap). Transcript annotation was based on the Ensembl genes database (release 77). To quantify gene expression levels, the number of reads mapped to the exons of each RefSeq gene was calculated. For older results using NGS pipeline version 3.16 and prior we used GRCH37 and Ensembl genes data base (release 67).

### Downstream analysis of bulk-RNA sequencing

Differential expression analysis was performed using DESeq2^63^. Signature scores were computed using the mean log-normalized expression of all genes in the signature and scaled using z-scoring. A two-sided student’s t-test was used to statistically test the difference in signatures between groups.

### Tissue processing and histology

Skin samples were formalin-fixed, paraffin embedded, sectioned (4-5 µm) and sections stained with Hematoxylin and Eosin or Alcian Blue. Slides were scanned on the Nanozoomer S360 platform (Hamamatsu) at 20x magnification for measurements.

### MVA ear infection

Mice were infected with Modified Vaccinia Virus Ankara (MVA; ATCC #VR-1508) as previously described^46,64^. Virus was propagated per ATCC guidelines in the BHK-21 cell line using standard protocols. Briefly, mice were anesthetized, and 10 µl of PBS containing 5 × 10⁶ PFU of MVA was topically applied to the ventral surface of the left ear pinna. The ear was then immediately scarified with 25 punctures using a sterile 29-gauge needle.

### Single Cell RNA-Sequencing

Infected ears and subcutaneous tumors were dissociated as described above. Cells were simultaneously stained and indexed with mouse TotalSeq-B hashtag antibodies (Biolegend; B0301-B0308) and a FACS antibody cocktail (EPCAM, CD45.2, TCRB, PDPN). Samples were washed twice with MACS buffer and pooled together into a single tube for further labeling with live-dead enrichment viability dyes (Calcein Violet and 7AAD). Cells were sorted for live cell fractions (stromal: Calcein Violet^+^, 7AAD^-^, CD45^-^, PDPN^+^; T cells: 7AAD^-^, CD45^+^ and TCRB^+^). Cells were counted using a haemocytometer and resuspended in MACS buffer for scRNA-Seq using the Chromium Single-Cell v2 3.1′ Chemistry Library, Gel Bead, Multiplex and Chip Kits (10x Genomics), according to the manufacturer’s protocol. A total of 10,000 cells were targeted per well.

### Library sequencing and single-cell data processing

Gene-expression libraries were sequenced on the NovaSeq platform (Illumina) with paired-end sequencing and dual indexing, whilst hashtag libraries were sequenced on the MiSeq platform (Illumina). A total of 26, 8 and 98 cycles were run for Read 1, i7 index and Read 2, respectively. FASTQ files from gene-expression and hashtag libraries were processed using the Cell Ranger Single Cell v.7.1.0 software (10x Genomics). In brief, scRNA-Seq data were processed using CellRanger count function (10x Genomics) and mapped to the mouse reference genome mm10 that was customized to contain the YFP sequence plasmid.

Filtered gene-cell count matrices that were output from the Cell Ranger count pipeline were imported into Seurat v.4.3.043^65–69^ using R v.3.6.1 and log-normalized using the *NormalizeData* function. Hashtag antibody count matrices were imported into Seurat and individual samples were demultiplexed using the demuxEM v0.1.744 package. Only cells determined as “Singlets” by demuxEM were included for analysis. Dimensional reduction was performed using the *SCTransform* workflow built within the Seurat package, which was utilized for downstream Principal Component Analysis (RunPCA) and clustering based on shared nearest neighbors (FindNeighbors) and leiden graph-based clustering using the original Louvain algorithm (FindClusters). A total of 30 principal components (default) were used as input for clustering steps and UMAP reduction (RunUMAP). Broad cell types were first annotated using the SingleR package^70^ with the ImmGen reference gene signature database. For CITE-Seq data processing, raw antibody-derived tag (ADT) counts were normalized using the DSB method^71^ to background signals from empty-droplets.

### Cluster annotation

For T cells at day 30 post MVA infection, a resolution of 0.4 was used and clusters were annotated for CD4 and CD8 T_RM_ that expressed combinations of canonical tissue-resident marker genes *Itgae* (CD103)*, Cd69, Cxcr6, Havcr2* and *Runx3,* and cell surface CD4 or CD8b, CD103, CXCR6 and CD69; *S1pr1^+^*and *Klf2^+^* circulating CD4 T cells (CD4_CIRC_)*; Mki67^+^ Top2a^+^* proliferating T cells; and a cluster of *Rora^+^, Nr3c1^+^, Il23r^+^* and *Igf1r^+^*innate lymphoid cells (ILCs) and gamma-delta T cells (ILCs_gdT).

For all cells in KPR^OVA^ tumors, a resolution of 1.2 was used and clusters were broadly annotated using ImmGen SingleR annotations as a guide for myeloid cells (*Fcer1g, Cd68, Adgre1*), NK and innate lymphoid cells (*Ncr1*), T cells (*Cd3e, Cd8a, Cd8b1*), B cells (*Jchain, Ms4a1, Cd79a*), endothelial cells (*Pecam1*), CAFs (*Col1a1, Pdpn, Lrrc15*), KPR^OVA^ tumor cells (*Krt8, Krt18*), and pericytes (*Myh11, Rgs5*).

For reclustering of CAFs from KPR^OVA^ tumors, a resolution of 0.4 was used and clusters were annotated for pan-tissue *Pi16^+^* fibroblasts (*Ly6c1, Cd34, Tnxb*), proliferative *Mki67^+^* fibroblasts, three clusters of inflammatory-like CAFs defined by *Cxcl12* (*Saa3, Mmp3, Sfrp2*), *Col15a1* (*Il34, Aspn, Il34*) and *Spp1* (*Sdc1, Cthrc1, Mmp9*), two *Lrrc15^+^* CAF clusters with myofibroblast-like (*Acta2, Tagln*) or matrix-remodeling (*Mmp10, Mmp13*) features, and a mitochondrial-high cluster likely reflecting apoptotic or metabolically stressed cells.

For reclustering of myeloid cells from KPR^OVA^ tumors, a resolution of 0.6 was selected and clusters were annotated for three tumor-associated macrophage (TAM) populations: *Spp1*^+^ inflammatory TAMs (*Itgam, Il1b*), *Apoe*^+^ lipid-associated TAMs (*Trem2, Mrc1, Sirpa, Adgre1, Cx3cr1*), and small cluster of osteoclast-like TAMs expressing Cathepsin K (*Ctsk*). Additional clusters included *Ccr2*^+^ monocytes, monocyte-derived TAMs (*Ccr2, Il1b*), monocyte-derived DCs (*Ccr2, Cd209a*), migratory DCs (*Ccr7*), plasmacytoid DCs (*Bst2, Siglech, Xcr1, Ly6c2*), neutrophils (*S100a8, S100a9*), mast cells (*Tpsb2*), and basophils (*Gata2*).

For CD8 T cells from KPR^OVA^ tumors, a resolution of 0.6 was selected and clusters were annotated for states that resembled stem-like cells (*Tcf7, Ly6c1*), IFN-responsive (*Ifit1, Ifit3)*, two cytotoxic effector states (*Gzmb*^+^ and *Gzmk*^+^), a transitional population, cycling cells (*Mki67, Top2a*), and a *Tox*^+^PD-1^+^ dysfunctional population enriched for proteotoxic stress response genes (*Hspe1, Hspa1b, Hsp90aa1, Bcl2l11*) (Fig. 7e).

Differential abundances between KPR^OVA^ tumors grown in WT and Floxed groups were computed using the Milo method^49^ and run using default parameters as recommended by the developers. Pseudo-bulk differential expression analysis was performed using DESeq2^63^ and pooling counts across all cells within each replicate. Gene signatures were scored across all cells using the AUCell method^72^. AUCell scores were calculated in only the top 5% of genes ranked by log-normalized expression, as recommended by the developers.

### SCimilarity and cross-tissue cell atlas analysis

Previously annotated human CD8 T_RM_ from lung^39^, breast cancer^14^, and ulcerative colitis^40^ studies were queried in the SCimilarity^38^ cell foundation model following the developers parameter recommendation. CD8 T_RM_ were defined by a SCimilarity distance metric cutoff (0.0627) and a core CD8 T_RM_ signature^10^ scored using AUCell^72^, defined against circulating blood CD8 and CD4 T cells (0.06). CD8 T_RM_ and all other cell type abundances from gut, lung and skin related datasets were extracted based on the number of cells divided by the total number of cells per sample in the SCimilarity database. Pearson correlations were computed on cell abundances in a pair-wise manner using the *cor* and *cor.test* functions in R. Similar Pearson correlation analyses on cell abundance were performed on healthy tissue and tumor cell atlas collated datasets from the Shi et al (2025) study^41^. Cell-cell communication predictions between fibroblasts and CD8 T_RM_ clusters from the Shi et al (2025) study^41^ were performed using the CellChat package^42^.

### Identification of cNMF programs

The cNMF method^73^ was used to identify biological gene programs across T cell compartments analyzed by scRNA-Seq. Feature selection was performed using only genes detected in at least 5% of cells and were identified as highly variable in Seurat (*FindVariableFeatures;* selection method = vst). This resulted in an input matrix for cNMF with a total of 1,000 and 2,000 genes for T cells from MVA infection and KPR^OVA^ scRNA-Seq experiments, respectively. cNMF was run with 100 iterations and with a number of components from 2 to 12. A k of 12 and 6 was selected as the preferred k considering stability and error. A threshold for euclidean distance of 0.1 was used to remove outliers. NMF program scores within cells were normalized to sum to 1. The top 100 weighted genes per program were used for gene ontology analysis with the Biological Processes sub-ontology database in the ClusterProfiler package^74–76^.

### In vivo tumour studies

Subcutaneous tumour experiments were performed as previously described^1^. The KPR PDAC tumor cell line (Genentech gCell) was maintained in DMEM containing 10% FBS and pen/strep antibiotics. KPR cells were first engineered to express the ovalbumin (OVA) model antigen using a piggybac transposase system (Transposagen Biolabs https://transposagenbio.com/) as previously described^77^. All constructs were synthesized by Genscript USA, Inc. The OVA expression cassette (pB_EF1_Ovalbumin) was preceded by an Elongation Factor 1 alpha (EF1α) promoter. Lipofectamine 3000 (Invitrogen) was used to transfect KPR cells with the piggybac plasmid along with a pBo transposase according to manufacturer protocol. Transfected cells were allowed to grow for 72 hours in DMEM, after which the cells were harvested and stained with the 25-D1.16 antibody (Biolegend, #141606) to identify KPR cells expressing the OVA-derived peptide SIINFEKL bound to H-2Kb of MHC class I. Single KPR cells with positive staining for 25-D1.16 were sorted by FACS into 96-well plates and maintained over 2-3 weeks at 37 C. Of all the expanded clones, the highest expressing KPR^OVA^ line determined by 25-D1.16 staining was selected for subsequent in vivo experiments.

For all mice, the right subcutaneous flank skin was shaved to remove hair the day prior to injection. KPR^OVA^ were grown in tissue culture in DMEM with 10% FCS. On the day of tumor implantation, cells were trypsinized, filtered, counted and resuspended in a 1:1 mixture of Hanks’s buffered saline solution and phenol-red-free Matrigel (Corning). Mice were anaesthetized using inhalatory anaesthesia during the tumor implantation procedure. Each mouse was subcutaneously inoculated in the right unilateral flank with 1 × 10^5^ KPR^OVA^ tumor cells in a total volume of 200 ul. Tumour volumes were measured and calculated 2-3 times per week using the following modified ellipsoid formula: ½ × (length × width^2^). Tumors that exceeded a volume of >1,000 mm^3^ were considered progressed and animals were removed from the study. Similarly, animals for which tumours ulcerated greater than 5 mm were removed from the study.

EMT6 tumor studies were performed in BALB/c mice. EMT6 cells were cultured in RPMI 1640 medium plus 2 mM l-glutamine with 10% fetal bovine serum (FBS; HyClone, Waltham, MA). For injections, cells were centrifuged, washed once with HBSS, counted, and resuspended in 50% HBSS and 50% Matrigel (BD Biosciences; San Jose, CA) at a concentration of 1 × 10^6^ cells/mL. A total of 1 × 10^5^ EMT6 cells were inoculated in the left mammary fat pad #5 of the mouse in 100 μL of HBSS: Matrigel (1:1). When the tumor reached a volume of 130–230 mm^3^, animals were distributed into treatment groups based on tumor volume and treated with isotype control antibodies (mouse IgG1 anti-gp120, 20 mg/kg first dose followed by 15 mg/kg thereafter), anti-PD-L1 (mouse IgG1 clone 6E11, 10 mg/kg first dose followed by 5 mg/kg thereafter), anti-TGF-β3 (2A10)^44^ or anti-TGF-β1^78,79^ (mouse IgG1, 10 mg/kg), or a combination of anti-PD-L1 with anti-TGF-β3 (2A10)^44^ or anti-TGF-β1^78,79^ (mouse IgG1, 10 mg/kg). Antibodies were administered three times a week for 21 days, the first dose intravenously, and subsequent doses intraperitoneally. Mice were euthanized immediately if tumor volume exceeded 2000 mm^3^.

## Data availability

Processed data, raw sequencing files and analysis code will be deposited to Open Science Framework (OSF) and a public sequencing data repository upon publication. No new algorithms were developed for this manuscript.

## Acknowledgements

We thank our colleagues in the FACS core at Genentech and the excellent support that they provided for flow sorting. We would like to thank Søren Warming and the rest of our talented colleagues in Molecular Biology for allele generation and genetic analysis. We thank our colleagues in the NGS core who provided support for sequencing. We thank members of the Turley, Muller and Krishnamurty laboratories for insightful discussions. We thank the Animal Resources department for their help with animal husbandry. This work was supported by Genentech.

## Author Contributions

S.Z.W. designed and performed experiments, analyzed and interpreted the experimental and bioinformatics data, and wrote the paper. R.S.L. help establish and perform viral experimental protocols. E.K.S. led the colon tissue dissociation experiments. H.B. led the pathology for the mouse models. A.C.V. and A.G. assisted with in vivo and imaging experiments. A.C. and Y.Y. provided insightful knowledge of TGFb biology and performed TGFb blockade in vivo experiments. T.S. provided insightful knowledge of TGFb3 biology. C.C. provided insightful experimental and biological knowledge of the colon. J.A.S. provided critical stromal and T cell knowledge throughout the project. A.T.K. provided critical feedback throughout the project and edited the paper. S.M. supervised the bioinformatics aspects of the project and edited the paper.

S.J.T. oversaw the project, interpreted the results and edited the paper.

## Supplementary Figures

**Extended Data Figure 1.**
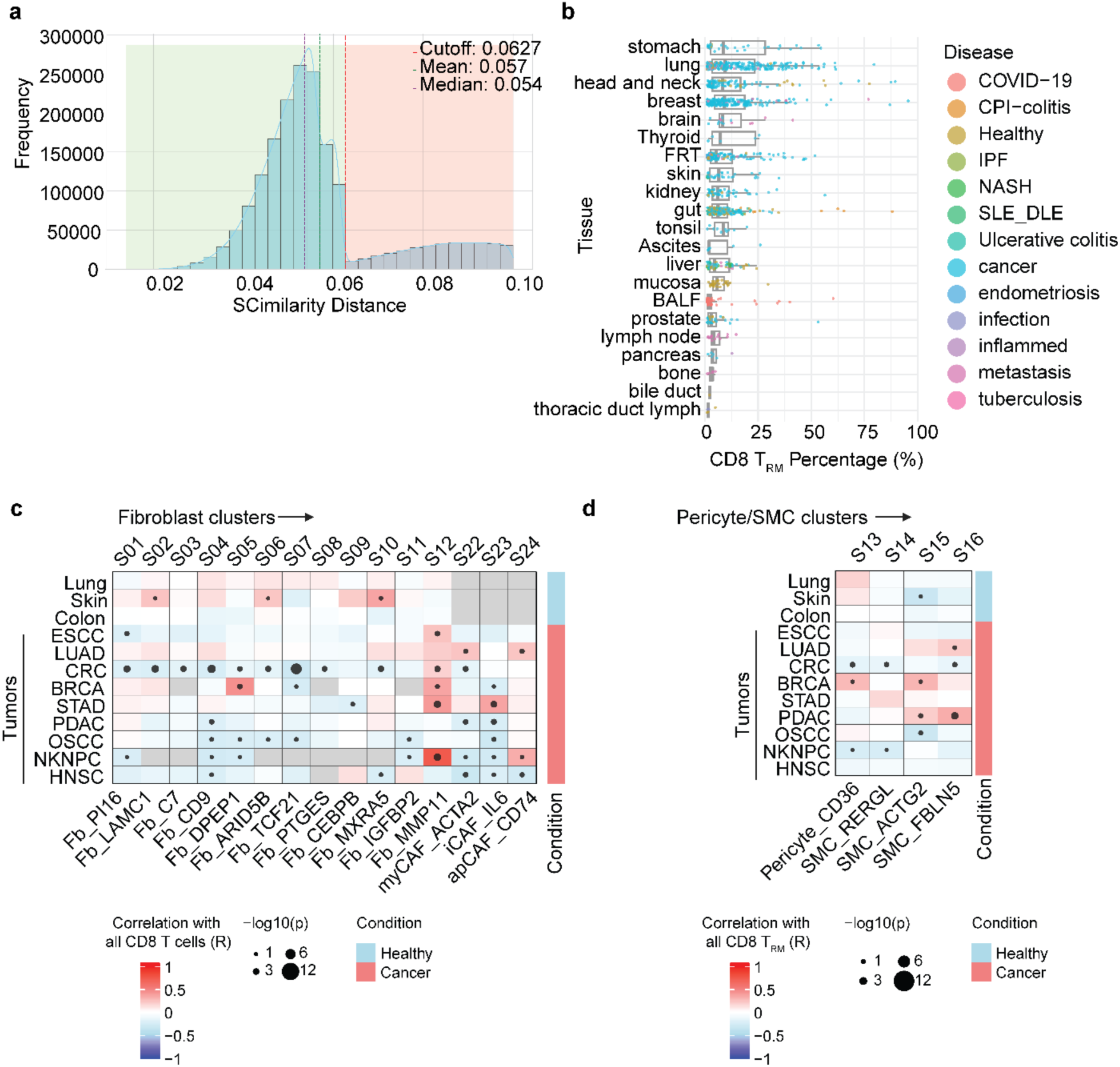
SCimilarity query of CD8 T_RM_ in human single cell datasets. **a**, Histograms of similarity scores as defined using SCimilarity. Threshold for filtering indicated by the red line. Higher SCimilarity scores indicate further distance from the query cells. **b**, Box plot summary of the frequency of CD8 T_RM_ detected across different tissue and disease types using SCimilarity. **c-d,** Pearson correlation analysis between independently annotated total CD8⁺ T cells and fibroblast clusters (c), and CD8 T_RM_ and pericytes/SMCs (d), in the cross-tissue atlas from Shi et al 2025^41^.

**Extended Data Figure 2.**
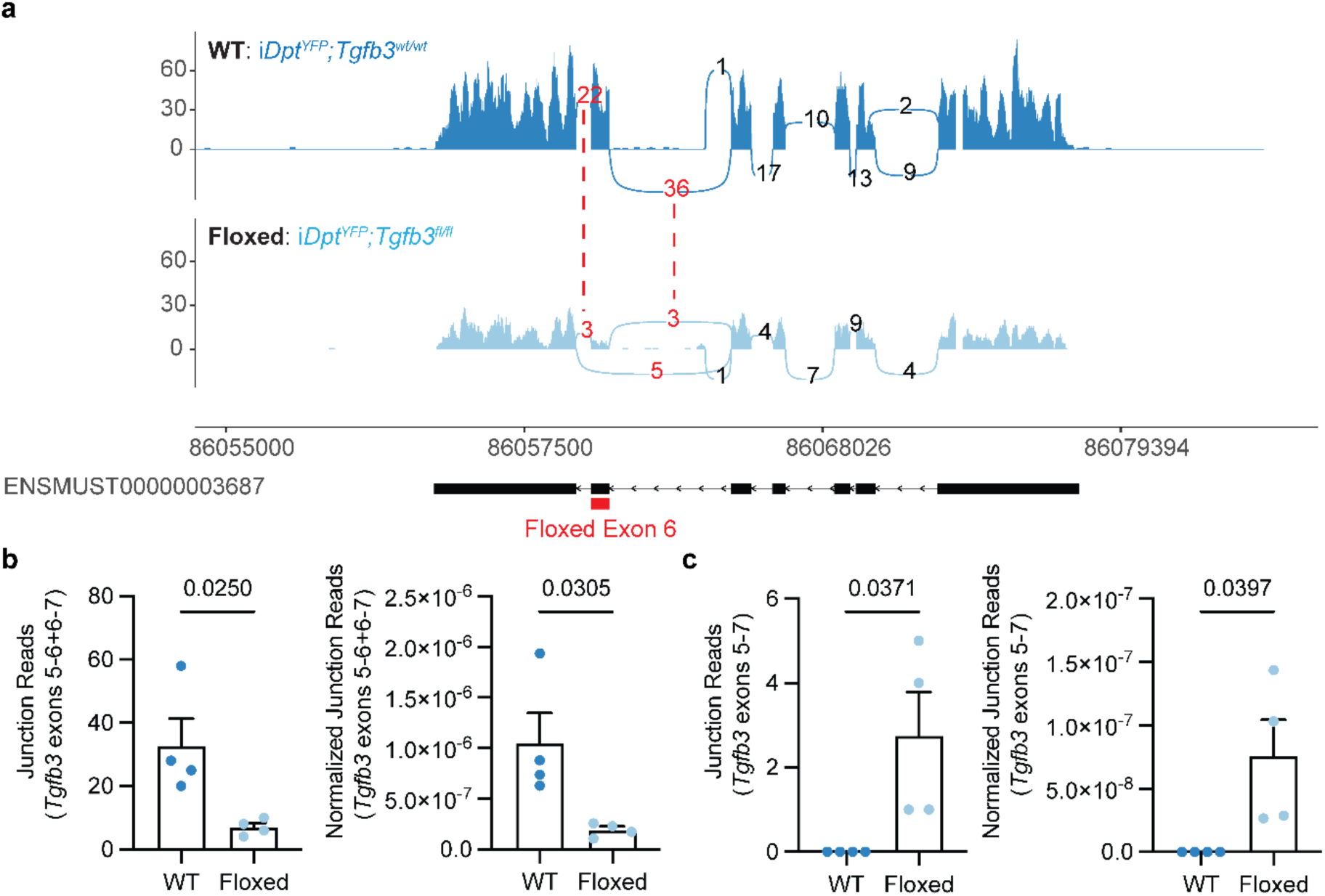
Deletion of *Tgfb3* exon 6 in *iDpt^YFP^Tgfb3^fl/fl^* mice. **a**, Representative ggsashimi plot showing the reduction of *Tgfb3* exon 6 junction reads in skin tissue profiled by bulk RNA sequencing in *iDpt^YFP^Tgfb3^fl/fl^*mice compared to *iDpt^YFP^Tgfb3^wt/wt^* control mice. Red lines and numbers indicate matched regions of *Tgfb3* exon 6. **b,** Quantification of exon 6 junction reads defined by reads spanning exons 5-6 and exons 6-7 between *iDpt^YFP^Tgfb3^fl/fl^*mice compared to *iDpt^YFP^Tgfb3^wt/wt^* control mice (n=4 per group). **c,** Quantification of exon 5-7 junction reads defined by reads that skip floxed exon 6 between *iDpt^YFP^Tgfb3^fl/fl^* mice compared to *iDpt^YFP^Tgfb3^wt/wt^* littermate control mice (n=4 per group).

**Extended Data Figure 3.**
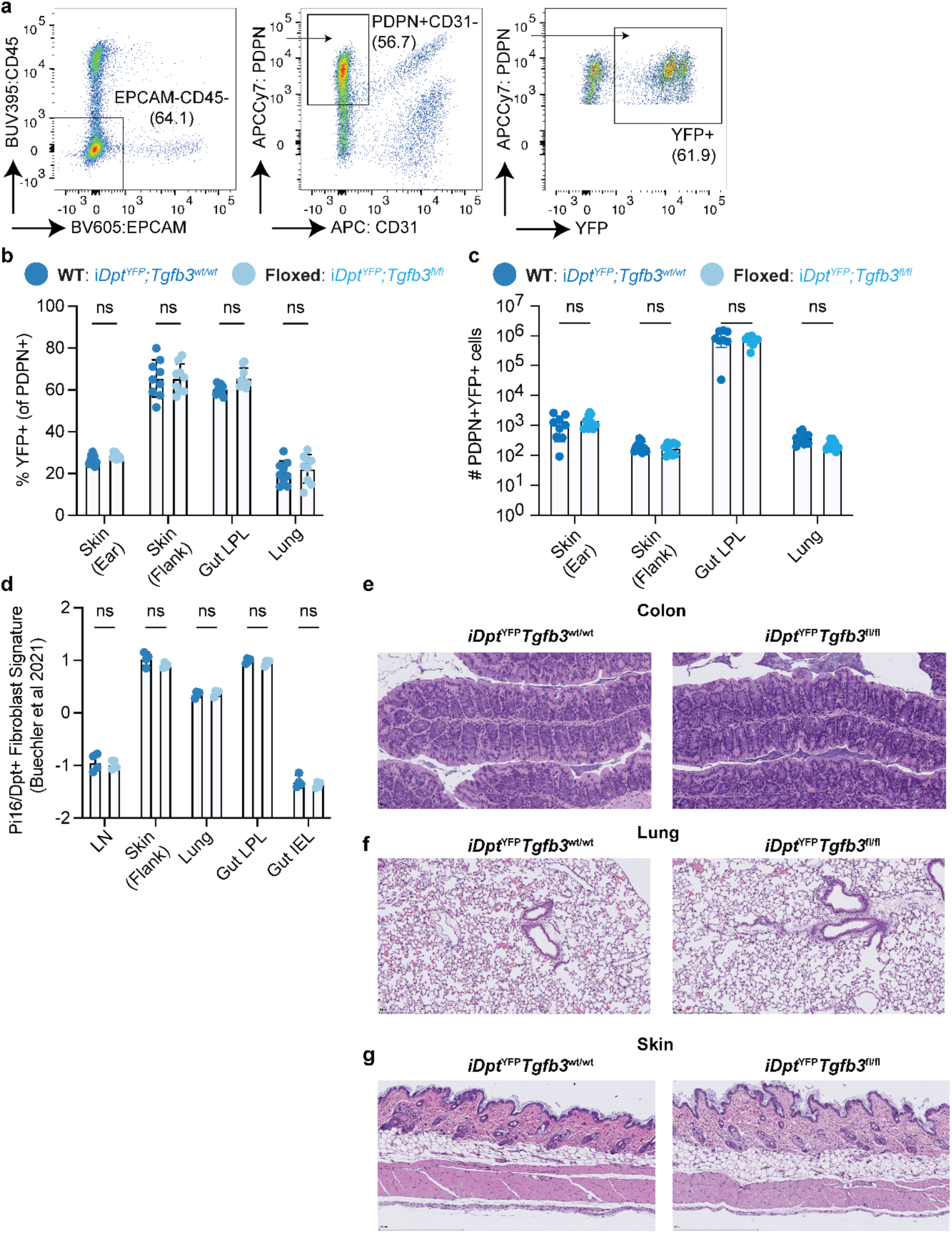
Fibroblasts and barrier tissue histology are unaltered in *iDpt^YFP^Tgfb3^fl/fl^* mice. **a**, Representative flow cytometry of *Dpt*+ fibroblasts in barrier tissues defined using Live CD45^-^EPCAM^-^ CD31^-^ PDPN^+^ and YFP as a reporter for *Dpt* expression. **b-c,** Quantification of the frequency (b) and total number (c) of YFP^+^ fibroblasts in barrier tissues of *iDpt^YFP^Tgfb3^fl/fl^* mice and controls (n=9 per group). Enumeration for skin and lung are represented as total cells per mg tissue, whereas colon is shown as total cells per colon. **d**, Scoring of a *Pi16/Dpt^+^* fibroblast gene signature across bulk RNA-seq profiling of lymph nodes and barrier tissues from *iDpt^YFP^Tgfb3^fl/fl^*mice and controls (n=4 per group). Signature from Buechler et al 2021^43^. Representative data from three independent experiments shown for b-c. **e-g,** Representative H&E images of colon (e), lung (f) and skin (g) histological analyses between *iDpt^YFP^Tgfb3^fl/fl^*mice and controls (n=5 per group). Statistical significance determined using a two-way ANOVA in b, c and d.

**Extended Data Figure 4.**
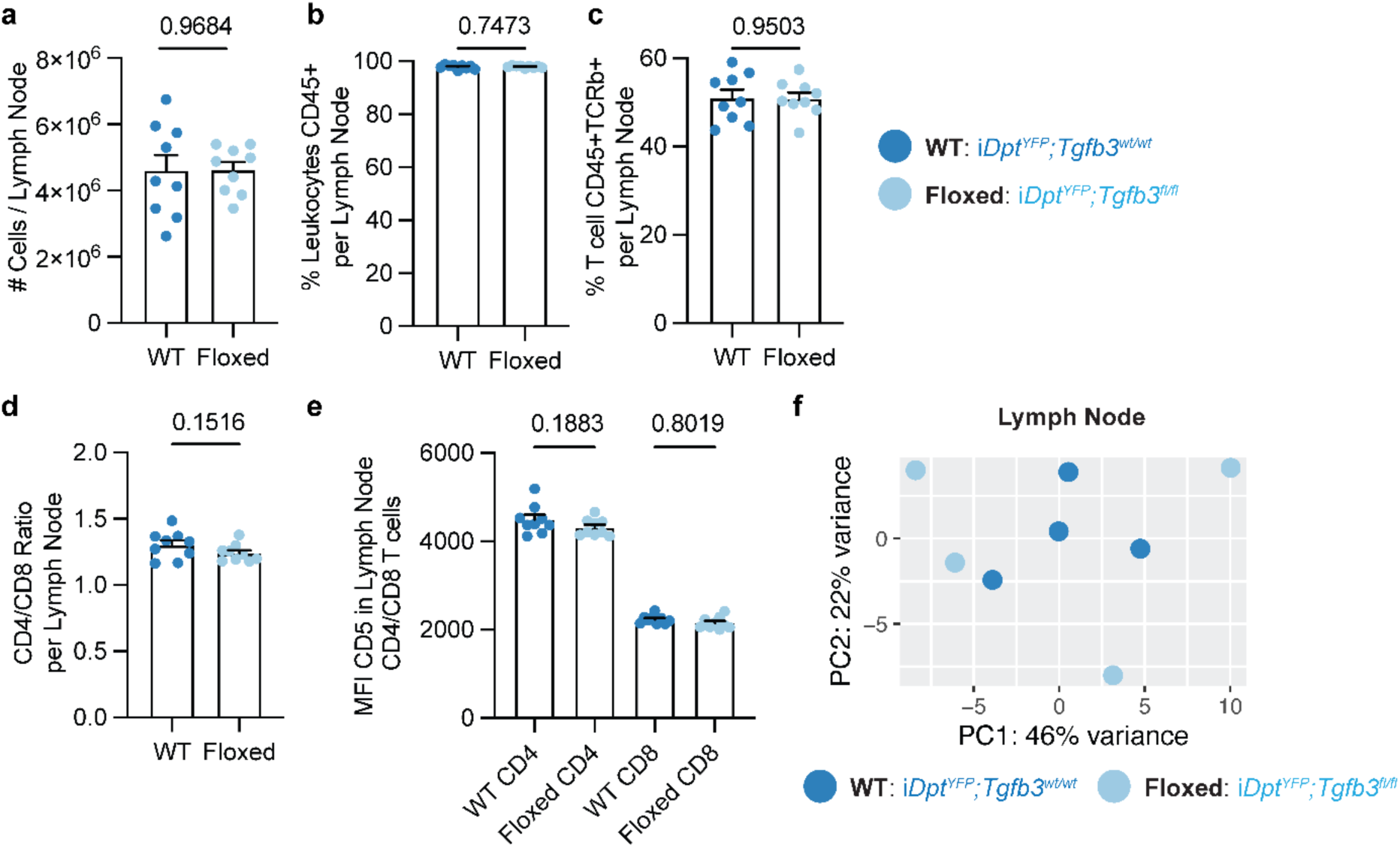
Lymph node immune compartments are unaltered in *iDpt^YFP^Tgfb3^fl/fl^* mice. **a-e**, Quantification of the total cellularity (a), frequency of CD45^+^ cells (b), frequency of CD45^+^ TCRb+ cells (c), ratio of CD4 and CD8 T cells (d) and median fluorescence intensity (MFI) of CD5 training in CD4 and CD8 T cells (e) between *iDpt^YFP^Tgfb3^fl/fl^* mice and controls (n=9 per group). **f,** PCA of lymph nodes sampled using bulk RNA sequencing from *iDpt^YFP^Tgfb3^fl/fl^* mice and controls (n=4 per group). Representative data from three independent experiments shown for a-e. Statistical significance determined using a two-sided unpaired t-test for a-d and two-way ANOVA in e.

**Extended Data Figure 5.**
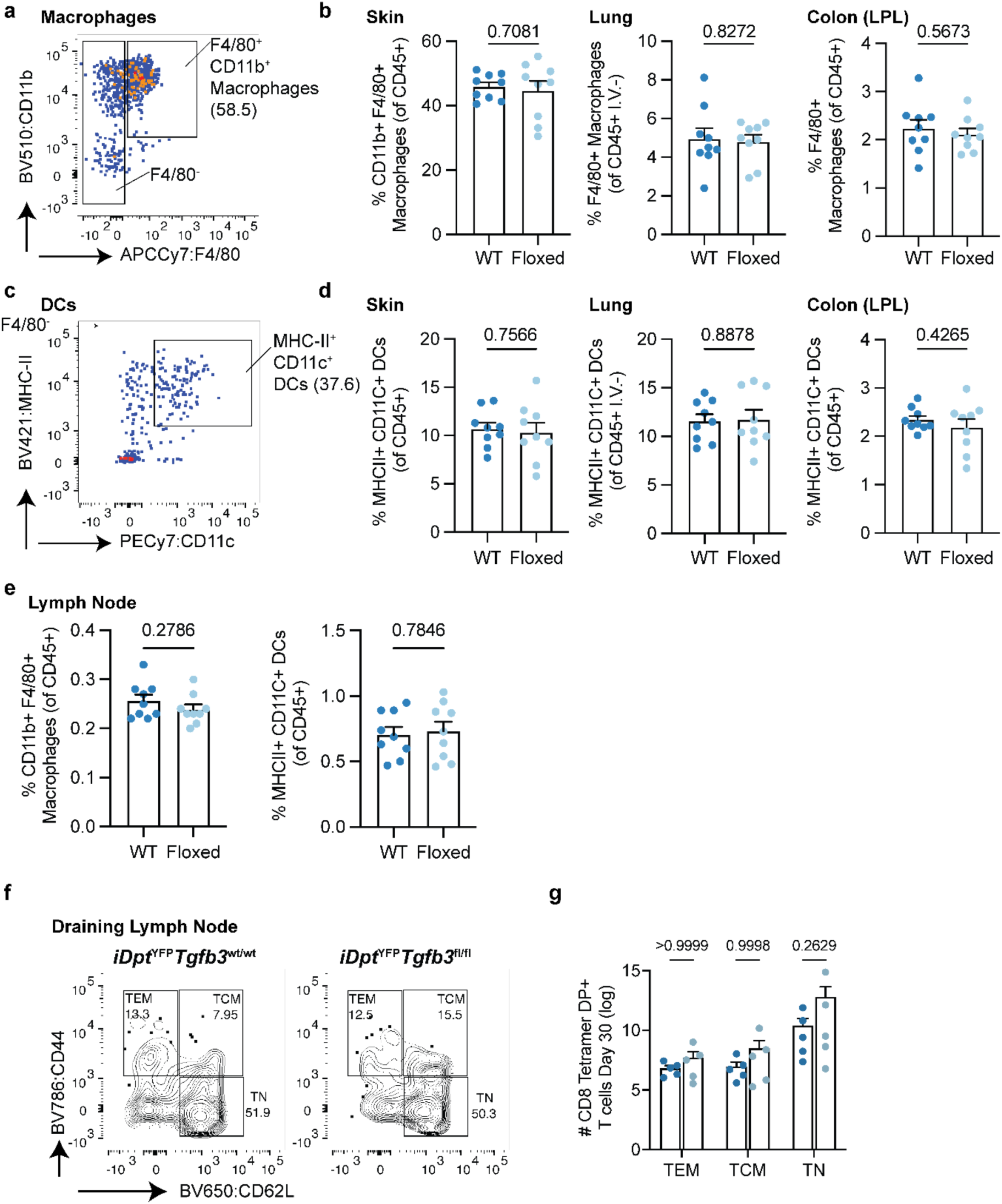
Myeloid cells and infected ear skin draining lymph nodes are unaltered in *iDpt^YFP^Tgfb3^fl/fl^*mice. **a-b**, Representative flow cytometry and quantification of CD11b^+^F4/80^+^ macrophages in barrier tissues from *iDpt^YFP^Tgfb3^fl/fl^* mice and controls (n=9 per group). Cells gated on Live CD45^+^. **c-d,** Representative flow cytometry and quantification of CD11c^+^ MHC-II^+^ dendritic cells in barrier tissues from *iDpt^YFP^Tgfb3^fl/fl^*mice and controls (n=9 per group). Cells gated on Live CD45^+^ CD11b^+^ F4/80^-^. **e**, Quantification of lymph node CD11b^+^F4/80^+^ macrophages and CD11c^+^ MHC-II^+^ dendritic cells in from *iDpt^YFP^Tgfb3^fl/fl^*mice and controls (n=9 per group). **f-g,** Representative flow cytometry and quantification of virus-specific CD44^HIGH^/ CD62L^LOW^ effector memory, CD44^HIGH^/ CD62L^HIGH^ central memory, and CD44^LOW^/ CD62L^HIGH^ naïve CD8 T cell populations in draining cervical lymph nodes at day 30 post MVA infection from *iDpt^YFP^Tgfb3^fl/fl^*mice and controls (n=5 per group). Representative data from three independent experiments shown for a-g. Statistical significance determined using a two-sided unpaired t-test for a-d and two-way ANOVA in e.

**Extended Data Figure 6.**
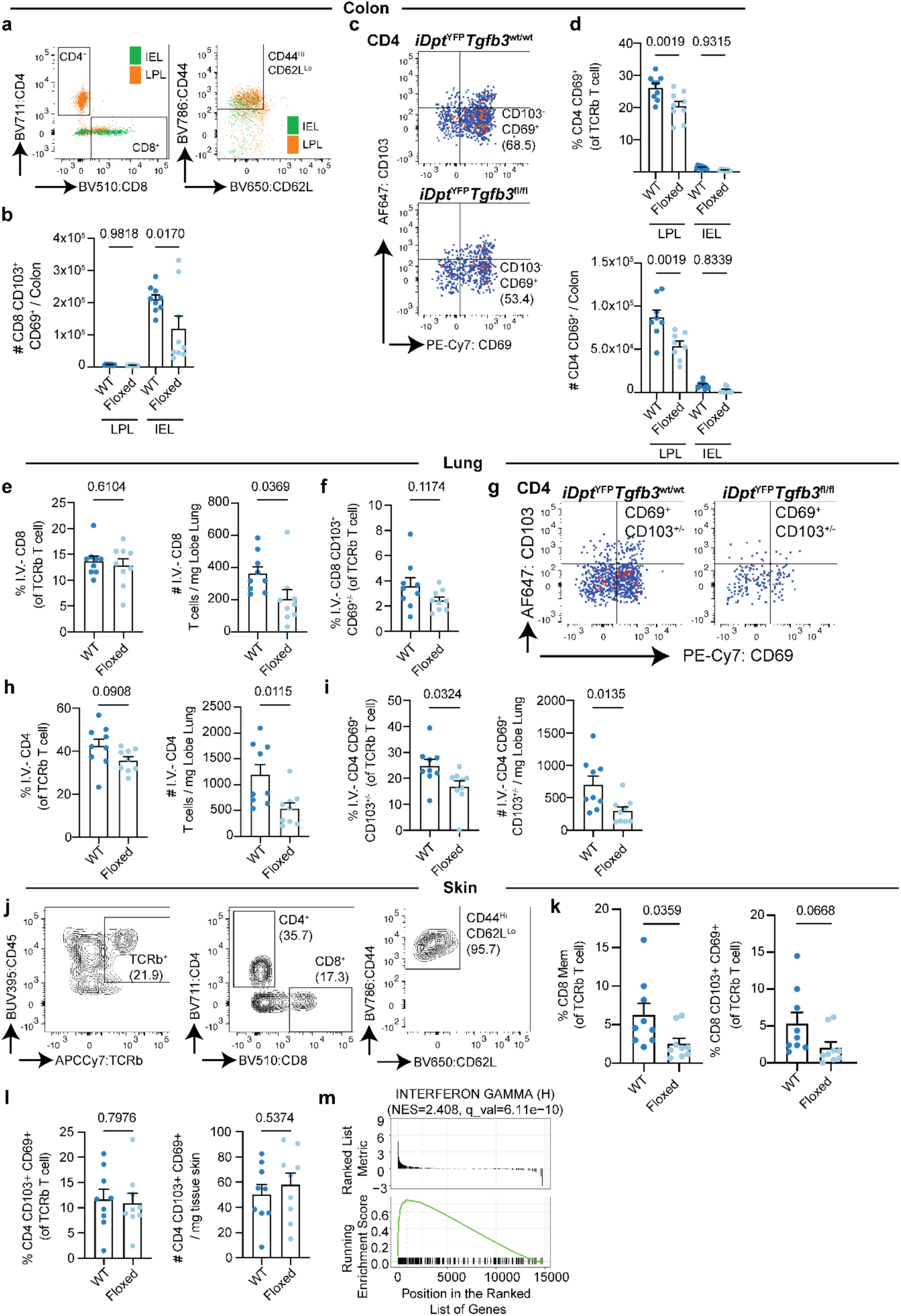
Barrier tissue CD4 and CD8 T_RM_ are altered in *iDpt^YFP^Tgfb3^fl/fl^* mice. **a**, Representative flow cytometry of CD4 and CD8 T cells in colon IEL and LPL compartments from *iDpt^YFP^Tgfb3^fl/fl^*mice and controls. Cells gated on Live CD45⁺ TCRβ⁺. **b,** Quantification of the total number of colon CD8 T_RM_ in *iDpt^YFP^Tgfb3^fl/fl^*mice and controls (n=9 per group). CD8 T_RM_ defined on CD44^HIGH^ CD62L^LOW^ CD103^+^ CD69^+^. **c-d,** Representative flow cytometry (c) and quantification (d) of colon CD44^HIGH^ CD62L^LOW^ CD69^+^ CD4 T cells from *iDpt^YFP^Tgfb3^fl/fl^*mice and controls (n=9 per group).. **e**, Quantification of the frequency and total number of CD8 T cells in the lungs of *iDpt^YFP^Tgfb3^fl/fl^*mice and controls (n=9 per group). Cells gated on Live CD45.2 I.V.^-^ CD45^+^ TCRβ^+^. **f,** Quantification of the frequency of lung CD8^+^ T_RM_ cells in *iDpt^YFP^Tgfb3^fl/fl^*mice compared to control animals. **g-i,** Representative flow cytometry (g) and quantification (h-i) of the frequency and total number of CD4 T cells (h) and CD44^HIGH^ CD62L^LOW^ CD69^+^ CD4 T cells (i) in lungs of *iDpt^YFP^Tgfb3^fl/fl^* mice and controls (n=9 per group). **j-k,** Representative flow cytometry (j) and quantification of CD44^HIGH^ CD62L^LOW^ CD8 T cells and CD8 T_RM_ in skin (k) of *iDpt^YFP^Tgfb3^fl/fl^* mice and controls (n=9 per group). **l**, Quantification of the frequency and total number of CD44^HIGH^ CD62L^LOW^ CD103^+^ CD69^+^ CD4 T cells in skin of *iDpt^YFP^Tgfb3^fl/fl^*mice and controls (n=9 per group). **m,** Gene set enrichment analysis (GSEA) of the Hallmark interferon response pathway, showing enrichment in *iDpt^YFP^Tgfb3^wt/wt^*control skin compared to *iDpt^YFP^Tgfb3^fl/fl^* skin. Representative data from three independent experiments shown for a-g. Statistical significance determined using a two-sided unpaired t-test for b, d, e, f, h, i, k and l.

**Extended Data Figure 7.**
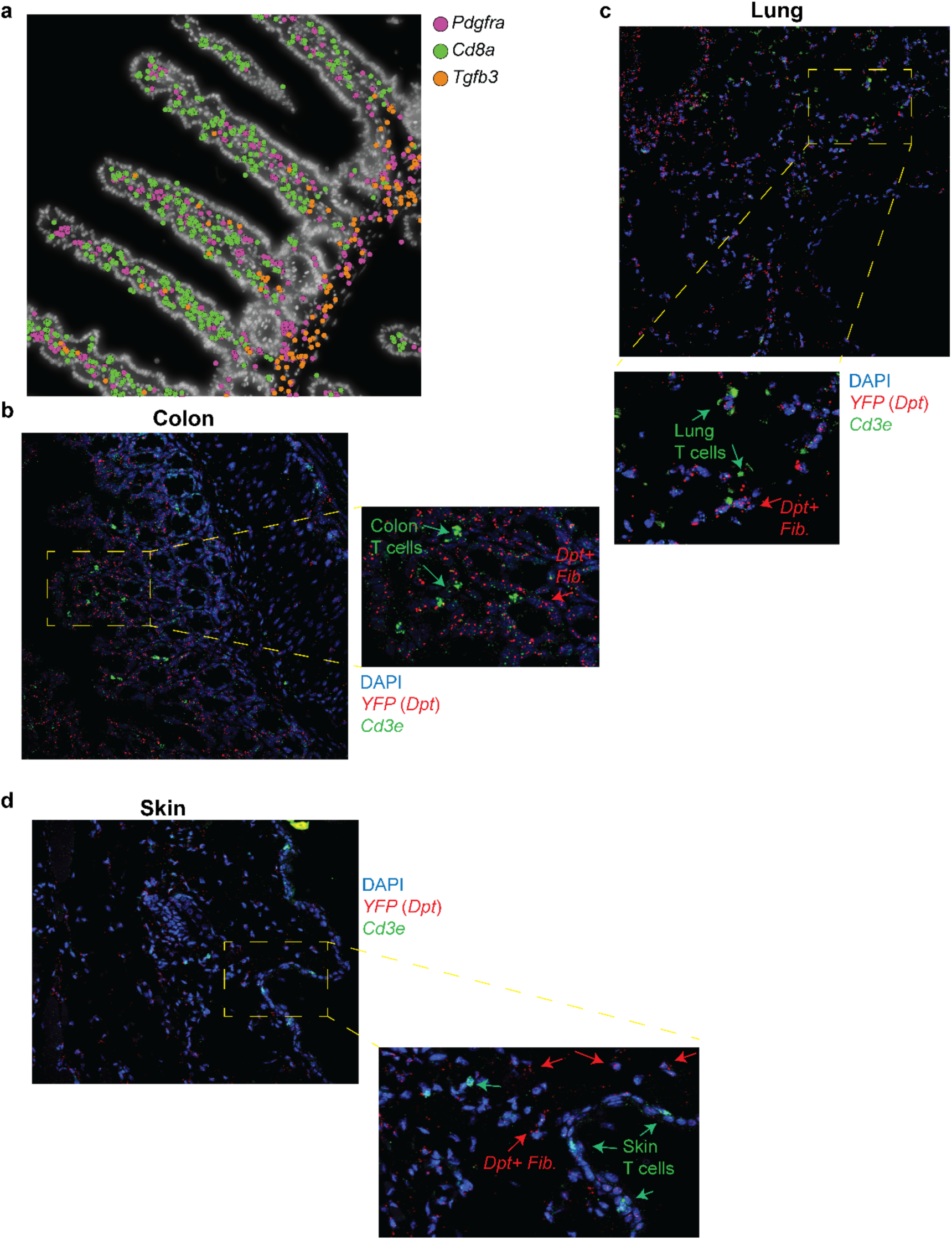
In-situ profiling of barrier tissue fibroblasts and T cells in *iDpt^YFP^Tgfb3^wt/wt^* mice. **a,** Spatial re-analysis of *Tgfb3*, *Cd8a* and *Pdgfra* expression in murine gut tissue from a published Xenium dataset (Reina-Campos et al.), illustrating fibroblast-restricted *Tgfb3* expression and proximity to CD8^+^ T cells. **b-d**, Representative images from in-situ hybridisation staining of *YFP* (red) and *Cd3e* (green) in colon (b), lung (c) and skin (d) of *iDpt^YFP^Tgfb3^wt/wt^*control mice. Demagnified images from Fig. 3 are shown. Representative images shown from 6 independent replicates per tissue.

**Extended Data Figure 8.**
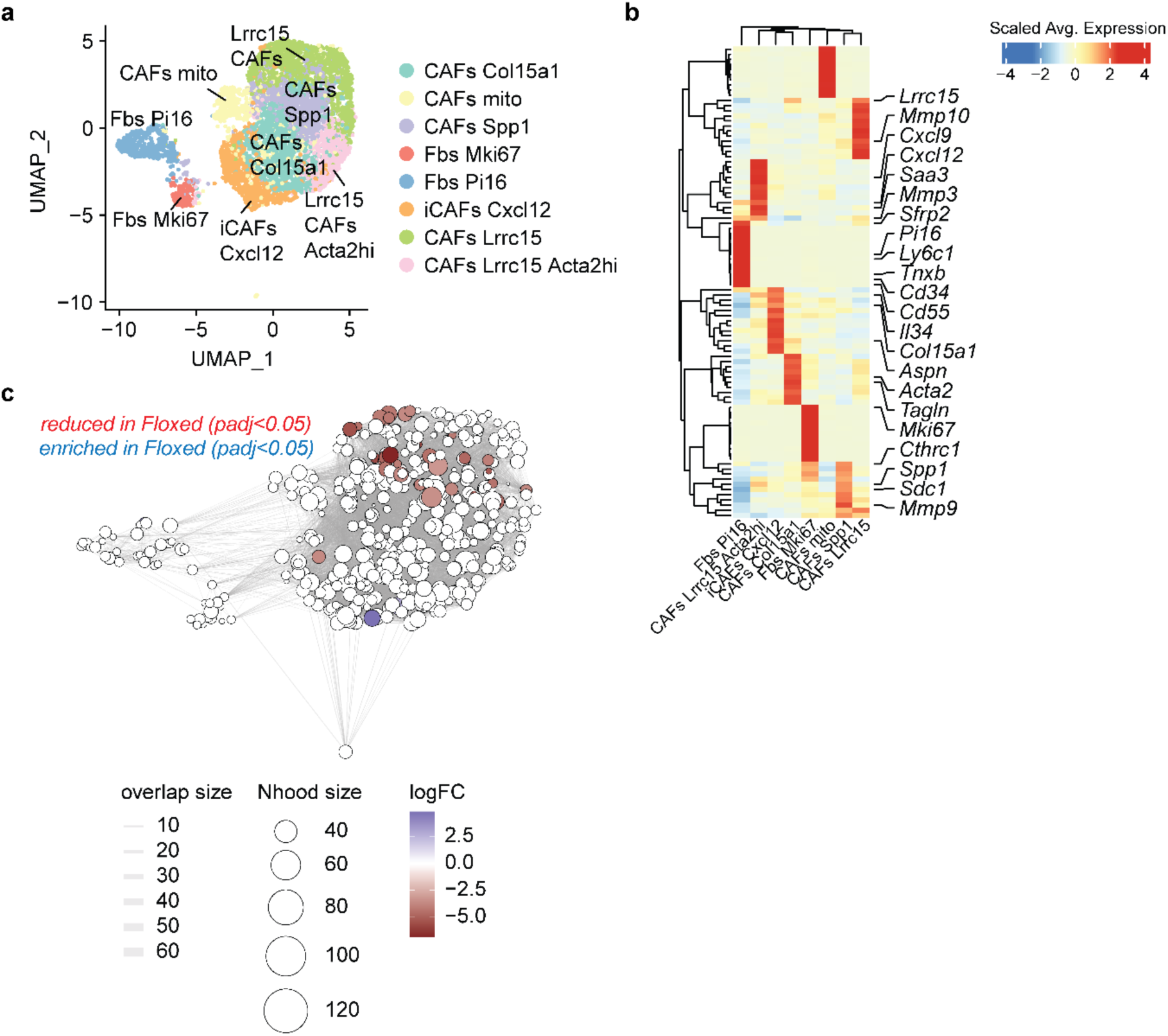
CAFs are largely unaltered transcriptionally in KPR^OVA^ tumors of *iDpt^YFP^Tgfb3^fl/fl^* mice. **a**, UMAP visualization of single-cell transcriptomic profiling of clustered cancer-associated fibroblasts (CAFs) isolated from KPR^OVA^ tumors grown in *iDpt^YFP^Tgfb3^fl/fl^* mice and controls. Reclustering revealed eight broad clusters including pan-tissue *Pi16^+^* fibroblasts (*Ly6c1, Cd34, Tnxb*), proliferative *Mki67^+^*fibroblasts, three clusters of inflammatory-like CAFs defined by *Cxcl12* (*Saa3, Mmp3, Sfrp2*), *Col15a1* (*Il34, Aspn, Il34*) and *Spp1* (*Sdc1, Cthrc1, Mmp9*), two *Lrrc15^+^*CAF clusters with myofibroblast-like (*Acta2, Tagln*) or matrix-remodeling (*Mmp10, Mmp13*) features, and a mitochondrial-high cluster likely reflecting apoptotic or metabolically stressed cells. **b**, Cluster-averaged heatmap of significantly differentially expressed genes across CAF clusters identified in a. **c,** Differential cell neighborhood analysis using Milo. A total of ∼8% of CAF neighborhoods were significantly reduced in *iDpt^YFP^Tgfb3^fl/fl^* mice compared to controls.

**Extended Data Figure 9.**
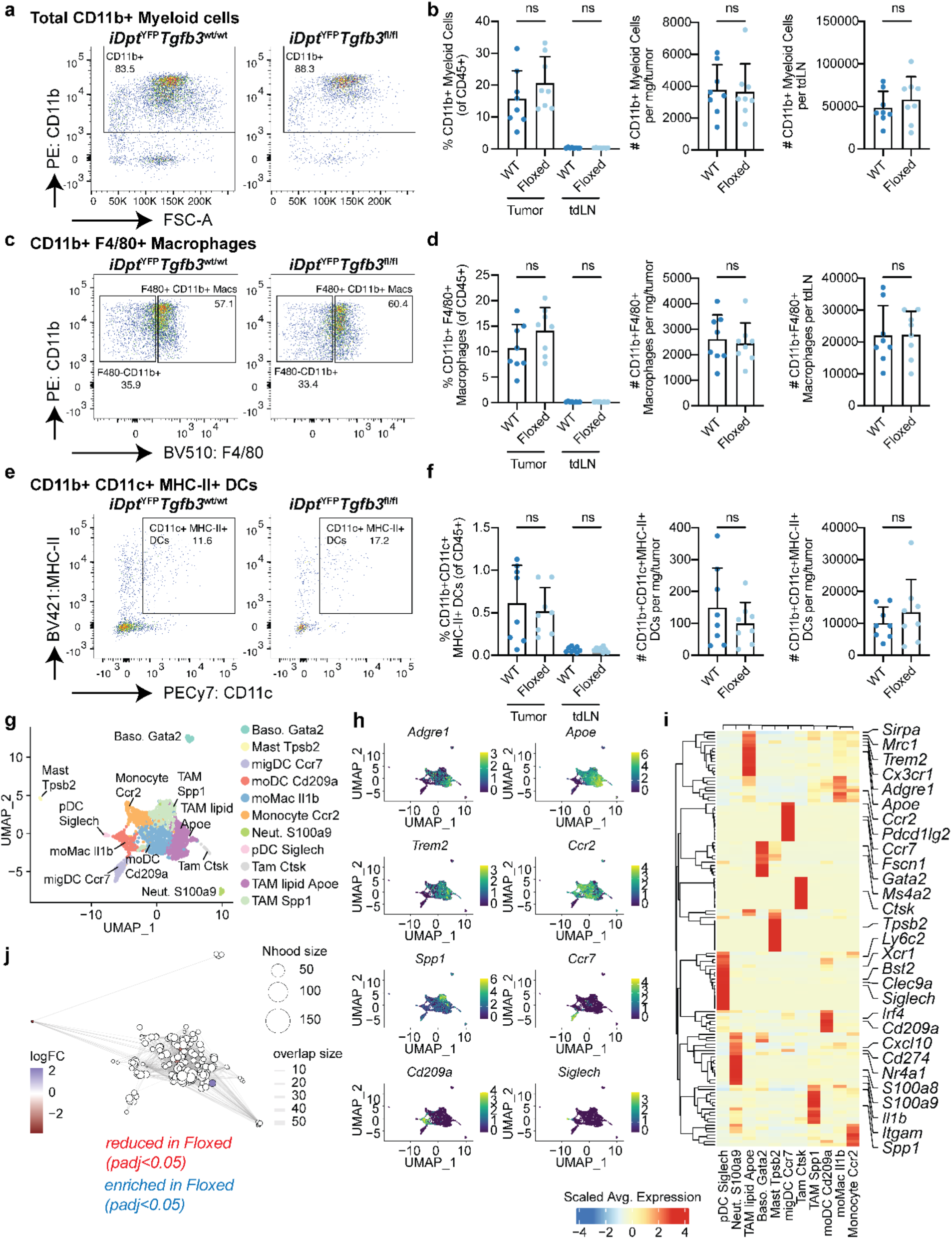
Myeloid cells are unaltered in KPR^OVA^ tumors of *iDpt^YFP^Tgfb3^fl/fl^* mice. **a-b**, Representative flow cytometry (a) and quantification (b) of the frequency and total number of CD11b+ myeloid cells from KPR^OVA^ tumors and tumor-draining lymph nodes (tdLNs) in *iDpt^YFP^Tgfb3^fl/fl^* mice and controls (n=7-8 per group). Cells gated on Live CD45^+^. **c-d**, Representative flow cytometry (c) and quantification (d) of the frequency and total number (d) of CD11b^+^F4/80^+^ macrophages from KPR^OVA^ tumors and tdLNs in *iDpt^YFP^Tgfb3^fl/fl^* mice and controls (n=7-8 per group). Cells gated on Live CD45^+^ CD11b^+^. **e-f**, Representative flow cytometry (e) and quantification (f) of the frequency and total number of CD11c^+^ MHC-II^+^ dendritic cells from KPR^OVA^ tumors and tdLNs in *iDpt^YFP^Tgfb3^fl/fl^*mice and controls (n=7-8 per group). Cells gated on Live CD45^+^ CD11b^+^ F4/80^-^. **g,** UMAP visualization of single-cell transcriptomic profiling of clustered myeloid cells isolated from KPR^OVA^ tumors grown in *iDpt^YFP^Tgfb3^fl/fl^* mice and controls. Eleven clusters include three tumor-associated macrophage (TAM) populations: *Spp1*^+^ inflammatory TAMs (*Itgam, Il1b*), *Apoe*^+^ lipid-associated TAMs (*Trem2, Mrc1, Sirpa, Adgre1, Cx3cr1*), and small cluster of osteoclast-like TAMs expressing Cathepsin K (*Ctsk*). Additional clusters included *Ccr2*^+^ monocytes, monocyte-derived TAMs (*Ccr2, Il1b*), monocyte-derived DCs (*Ccr2, Cd209a*), migratory DCs (*Ccr7*), plasmacytoid DCs (*Bst2, Siglech, Xcr1, Ly6c2*), neutrophils (*S100a8, S100a9*), mast cells (*Tpsb2*), and basophils (*Gata2*). **j,** Differential cell neighborhood analysis using Milo. A total of ∼3% of CAF neighborhoods were significantly reduced in *iDpt^YFP^Tgfb3^fl/fl^*mice compared to controls. **h-i**, Selected marker gene feature plots (h) cluster-averaged heatmap (i) of significantly differentially expressed genes across myeloid clusters. Representative data from three independent experiments shown for a-f. Statistical significance determined using two way ANOVA (left) and two-sided unpaired t-test (right) for b, d and f.

**Extended Data Figure 10.**
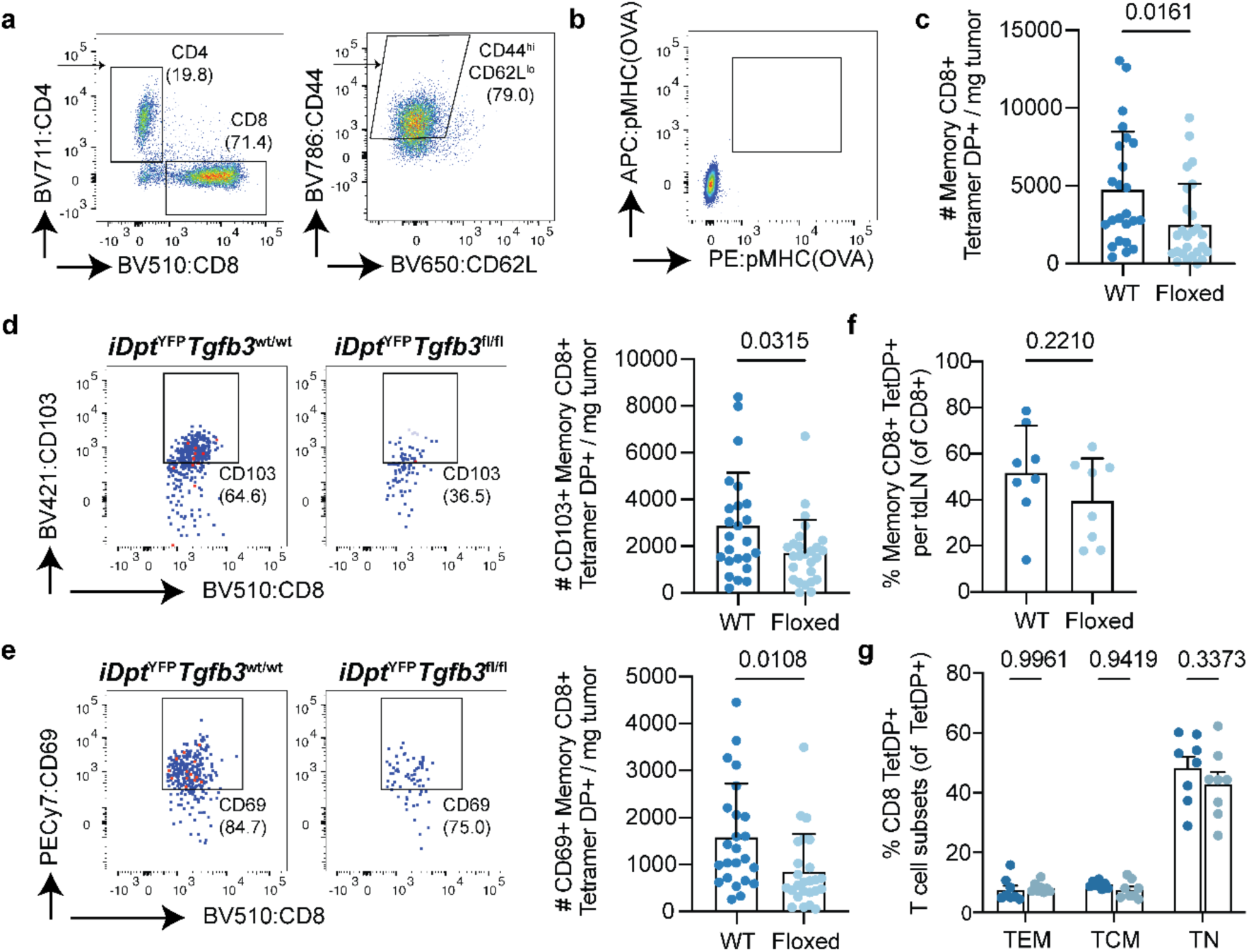
Antigen-specific CD8 T cells are reduced in KPR^OVA^ tumors of *iDpt^YFP^Tgfb3^fl/fl^* mice. **a-b**, Representative flow cytometry (a) of CD44^HIGH^ CD62L^LOW^ CD8 T cell gating in tumors and control tetramer negative tissue from tumor free mice (b). Staining of tetramer CD8 T cells tracked using H-2K(b) tetramers loaded with OVA peptide (SIINFEKL) on two different fluorophores. **c**, quantification of the total number of tumor-specific CD8 T cells in *iDpt^YFP^Tgfb3^fl/fl^* mice and controls (WT n=24, Floxed n=26).Tetramer^DP+^ cells gated on Live CD45^+^ YFP^-^ CD4^-^ CD8^+^ CD44^HIGH^ CD62L^LOW^. **d-e,** Representative flow cytometry and quantification of the total number of CD103*^+^* (d) and CD69*^+^* (e) tumor-specific CD8 T cells per mg of tumor tissue in *iDpt^YFP^Tgfb3^fl/fl^* mice and controls. **f-g,** Quantification of the total number of Tetramer^DP+^ CD8 T cells subsets in tumor-draining lymph nodes (tdLNs). Data pooled from three independent experiments shown for e, d and e. Statistical significance determined using two-sided unpaired t-test (right) for c, d, e and f, and two way ANOVA in g.

